# STREAM: Single-cell Trajectories Reconstruction, Exploration And Mapping of omics data

**DOI:** 10.1101/302554

**Authors:** Huidong Chen, Luca Albergante, Jonathan Y Hsu, Caleb A Lareau, Giosue` Lo Bosco, Jihong Guan, Shuigeng Zhou, Alexander N Gorban, Daniel E Bauer, Martin J Aryee, David M Langenau, Andrei Zinovyev, Jason D Buenrostro, Guo-Cheng Yuan, Luca Pinello

**Affiliations:** Molecular Pathology Unit & Cancer Center, Massachusetts General Hospital Research Institute and Harvard Medical School, Boston, MA 02114, USA; Department of Blostatlstlcs and Computational Biology, Dana-Farber Cancer Institute, Boston, MA 02215, USA; Department of Blostatlstlcs, Harvard T.H. Chan School of Public Health, Boston, MA 02215, USA; Department of Computer Science and Technology, Tongjl University, Shanghai 201804, China; Institut Curie, PSL Research University, F-75005 Paris, France; NSERM, U900, F-75005 Paris, France; MINES ParisTech, PSL Research University, CBIO-Centre for Computational Biology, F-75006 Paris, France; Department of Biological Engineering, Massachusetts Institute of Technology, Cambridge, MA, USA.; Broad Institute of MIT and Harvard, Cambridge, MA 02142, USA; Department of Mathematics and Computer Science, University of Palermo, Palermo 90123, ITALY; Department of Sciences for technological Innovation, Euro-Mediterranean Institute of Science and Technology, Palermo 90139, ITALY; Shanghai Key Lab of Intelligent Information Processing, and School of Computer Science, Fudan University, Shanghai 200433, China; Department of Mathematics, University of Leicester, University Road, Leicester LEI 7RH, UK; Lobachevsky University, Nizhni Novgorod, Russia; Division of Hematology/Oncology, Boston Children’s Hospital, & Department of Pediatric Oncology, Dana-Farber Cancer Institute Boston, Massachusetts, USA.; Harvard Society of Fellows, Harvard University, Cambridge, MA 02138, USA; Harvard Stem Cell Institute, Cambridge, MA 02138, USA

## Abstract

Single-cell transcriptomic assays have enabled the de novo reconstruction of lineage differentiation trajectories, along with the characterization of cellular heterogeneity and state transitions. Several methods have been developed for reconstructing developmental trajectories from single-cell transcriptomic data, but efforts on analyzing single-cell epigenomic data and on trajectory visualization remain limited. Here we present STREAM, an interactive pipeline capable of disentangling and visualizing complex branching trajectories from both single-cell transcriptomic and epigenomic data.

## Main text

STREAM (Single-cell Trajectories Reconstruction, Exploration And Mapping) can accurately recover complex developmental trajectories and provide informative and intuitive visualizations to highlight important genes that define subpopulations and cell types. STREAM reliably reconstructs trajectories and pseudotime (the distance from the start of a developmental trajectory) when multiple branching points are present, assumes no prior knowledge about the structure or the number of trajectories, and does not require extensive bioinformatics knowledge thanks to a user-friendly and interactive web interface. Additionally, STREAM has four innovations compared to other existing methods: 1) a novel density-level trajectory visualization useful to study subpopulation composition and cell-fate genes along branching trajectories, 2) a documented end-to-end pipeline to reconstruct trajectories from chromatin-accessibility data, 3) the first interactive database focused on single-cell trajectory visualization for several published studies, and 4) a trajectory mapping procedure to readily map new cells to precomputed structures without pooling data and recomputing trajectories. This last innovation allows facile analysis of data from genetic perturbation studies or to assign diseased/stimulated cells to a normal/resting developmental hierarchy. STREAM has been extensively tested using several published datasets from different organisms (zebrafish, mouse, human) and single-cell technologies (qPCR, scRNA-seq, scATAC-seq). It also has been compared to 10 other methods on both synthetic and real datasets.

STREAM takes as input a single-cell gene expression or epigenomic profile matrix and approximates the data in three or more dimensions with a structure called the principal graph, a set of curves that naturally describe the cells’ pseudotime, trajectories and branching points **(Fig. 1a)**. STREAM first identifies informative features such as variable genes or top principal components. Using these features, cells are then projected to a lower dimensional space using a non-linear dimensionality reduction method called Modified Locally Linear Embedding (MLLE), which preserves distances within local neighborhoods. In the MLLE embedding, STREAM infers cellular trajectories using a novel **El**astic **P**r**i**ncipal **Graph** implementation called ElPiGraph^1^. ElPiGraph is a completely redesigned algorithm for elastic principal graph optimization introducing the elastic matrix Laplacian, trimmed mean square error, explicit control for topological complexity and scalability to millions of points. In STREAM, ElPiGraph is integrated with a heuristic graph structure seeding and several graph grammars rules optimized for single-cell data. In contrast to the majority of existing methods, ElPiGraph does not rely on kNN graphs or minimum spanning trees. ElPiGraph is very robust to background noise, does not require pre-clustering, can work in multidimensional space, and is able to manage large-scale datasets on an ordinary laptop **(Online Methods)**.

**Figure 1.**
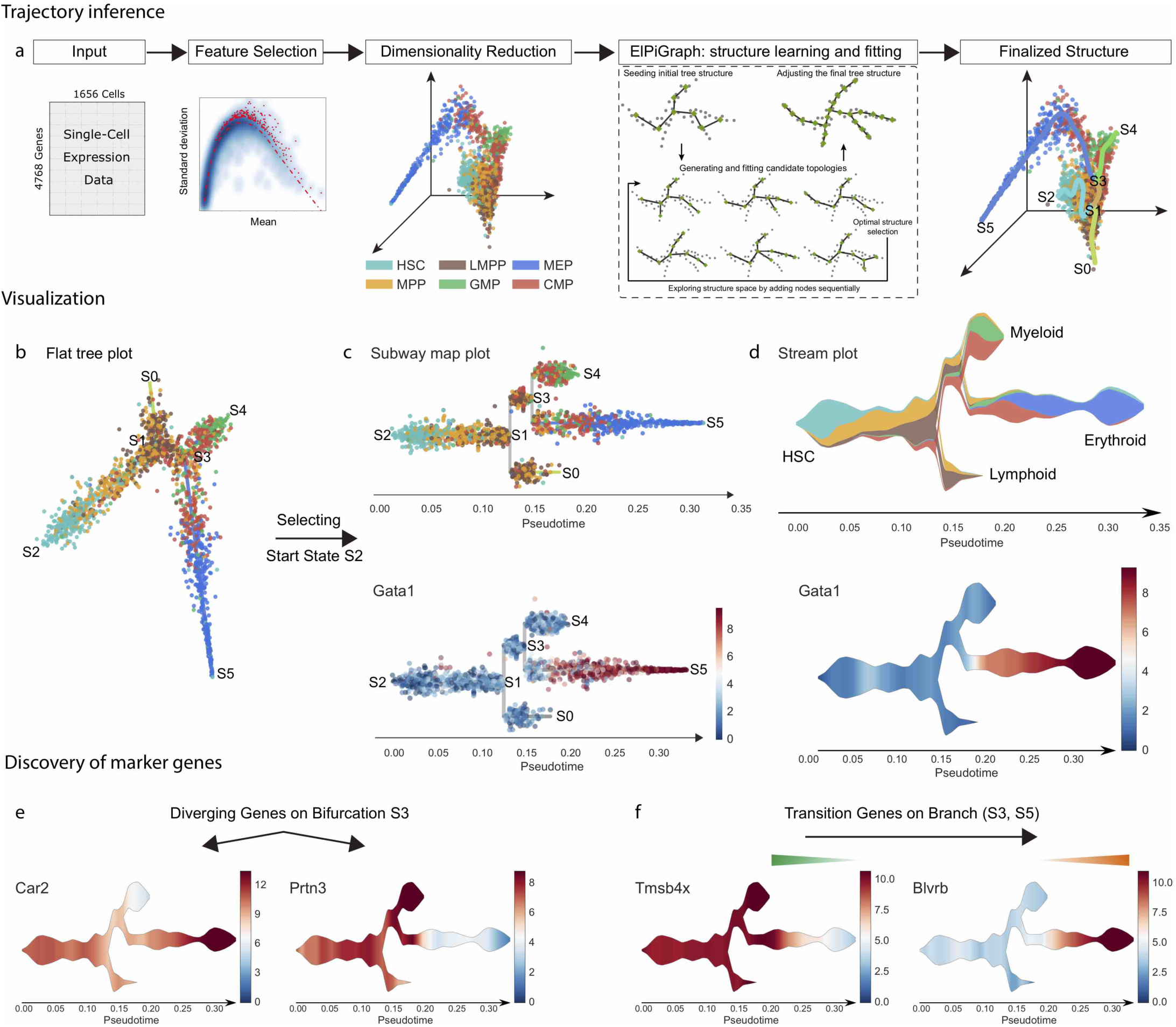
STREAM pipeline overview on single-cell RNA-seq data from the mouse hematopoietic system. **(a)** STREAM trajectory inference. Starting with a single-cell gene expression matrix, STREAM performs three main steps: selection of informative genes, dimensionality reduction, and simultaneous tree structure learning and fitting by ElPiGraph. The optimal structure is selected based on the *elastic energy* minimization among a set of candidate structures that are constructed every time a tree node is added. The final tree is interpreted as a set of connected curves representing different trajectories. **(b-d)** STREAM visualization of inferred branching points, trajectories and expression of key genes at both single-cell level and density level. **(b)** *Flat tree plot*, branches are represented as straight lines and each circle represents a single cell. The lengths of the branches and the distances between cells and their assigned branches are preserved from the space where trajectories were inferred. **(c)** *Subway map plot*, after selecting an initial state in the flat tree plot, the tree is re-ordered to facilitate visualization. Each cell is colored by a cell label, if provided (top), or based on the expression of a gene of interest (bottom). **(d)** *Stream plot*, a novel and intuitive visualization to show cell density along different trajectories: at a given pseudotime, the width of each branch is proportional to the total number of cells (top). Stream plots can also visualize the expression of a gene of interest (bottom). **(e-f)** STREAM detection of marker genes. **(e)** STREAM automatically discovers important marker genes for each branch. Left, identification of differentially expressed genes between bifurcating branches. **(f)** Identification of transition genes (expression values correlate with pseudotime) along one specific branch. Top two detected differentially expressed genes (*Car2* and *Prtn3)* and transition genes (*Tmsb4x* and *Blvrb)* are shown respectively with stream plots.

To illustrate STREAM, we first reanalyzed scRNA-seq data from Nestorowa et al. 2016^2^, which sorted and profiled 1,656 single mouse hematopoietic stem and progenitor cells. Starting from the hematopoietic stem cells (HSCs), STREAM accurately recapitulates known bifurcation events in lymphoid, myeloid and erythroid lineages and positions the multipotent progenitors before the first bifurcation event. To facilitate the exploration of the inferred structure, STREAM includes *a flat tree plot that* intuitively represents trajectories as linear segments in a 2D plane. In this representation, the lengths of tree branches are preserved from the MLLE embedding **(Fig. 1b)**. In addition, cells are projected onto the tree according to their pseudotime locations and the distances from their assigned branches. If the process under study has a natural starting point (for example a known origin in a developmental hierarchy), the user can specify a root node. This allows easy re-organization of the tree using a breadth-first search to obtain a *subway map plot* that better represents pseudotime progression from a selected starting node **(Fig. 1c)**. Although these visualizations capture trajectories and branching points, they are not informative on the density and composition of cell types along pseudotime, a common challenge when modeling large datasets. To solve this problem, we propose a novel trajectory visualization method called the *stream plot.* This compact representation summarizes cellular developmental trajectories, user-defined annotations, branching points, cell density, and gene expression patterns **(Fig. 1d)**. Density information, an aspect overlooked by other methods, is very important to track how the composition of subpopulations changes along a trajectory or gets partitioned around branching events. Additionally, STREAM detects potential marker genes of different types: *diverging genes,* i.e. genes important in defining branching points that are differentially expressed between diverging branches, and *transition genes*, i.e. genes for which the expression correlates with the cell pseudotime on a given branch. The expression patterns of the discovered genes can then be visualized using either subway maps or stream plots **(Fig. 1e-f, Supplementary Fig. 1-2, Supplementary Note 1)**.

STREAM is the only trajectory inference method that explicitly implements a *mapping* procedure, which allows reusing a previously inferred principal graph as reference to map new cells not included in the original fitting procedure. This avoids pooling old and new cells and re-computing trajectories from scratch, a computationally-intensive operation that also perturbs the original structure and complicates the interpretation of the pseudotime. This feature is particularly helpful when studying genetic perturbation data or exploring unlabeled data. To show the utility of the mapping feature, we applied STREAM to scRNA-seq data from *Olsson et al^3^*. This study focused on the mouse hematopoietic system, specifically on the consequences of cell determination within the granulocyte-monocyte progenitors (GMP) population after the transcription factors Gfi1 and/or Irf8 are knocked out. STREAM recovers the correct trajectories for the wild-type cells and, using the mapping feature, also predicts and effectively visualizes the consequences of the genetic perturbation as validated in the original study **(Supplementary Fig. 3-4, Supplementary Note 2)**.

To test the robustness and scalability of STREAM, we next explored data derived from different platforms and organisms. We used two recently published zebrafish datasets obtained with single-cell qPCR^4^ and inDrop^5^ (profiling ∼10000 cells) assays. These data provided the first comprehensive model of the zebrafish hematopoiesis system without biases introduced by FACS sorting subpopulations. Our analyses successfully recovered developmental trajectories at unprecedented resolution compared to previous analysis **(Supplementary Fig. 5-6, Supplementary Note 3-4)**.

We next systematically compared STREAM with 10 other state of the art methods for pseudotime inference on three different datasets^6-14^. First, we assessed the quality of the topology using a previously proposed synthetic dataset^14^. Second, we assessed the pseudotime accuracy using known marker genes on scRNA-seq data for myoblast differentiation, a classic dataset to compare trajectory inference methods^15^. Finally, we quantitatively compared the number and quality of trajectories in terms of precision and recall for known marker genes that completely diverge during development. In all the comparisons, STREAM consistently outperforms the other methods in inferring the correct topology, provides smooth pseudotime for myoblast differentiation and reconstructs the most balanced branching structure (avoiding under/over branching) in terms of precision and recall (F1-score) among the methods **(Supplementary Figs. 7-14, Supplementary Note 5)**.

Importantly, we extended STREAM to infer trajectories from human single-cell epigenomic data. This task is particularly challenging since the number of chromatin peaks (∼450,000 peaks across hematopoiesis) far exceeds the number of genes and the accessibility at each peak is sparse, often containing only 0, 1, or 2 reads. Additionally, trajectory reconstruction based on scATAC-seq human data is more difficult than the recently obtained trajectories in non-mammalian organisms^16^ with much smaller genomes. STREAM is able to perform pseudotime ordering on human cell chromatin-accessibility data without relying on accessibility of known transcription factor binding sites^17^ or *a priori* knowledge of sampling time^18^, hence providing a truly unbiased approach. STREAM in fact uses an unbiased set of DNA sequence features (7-mers), scoring each cell with chromVAR^19^ based on its accessibility deviations across cells **(Fig. 2a)**.

**Figure 2.**
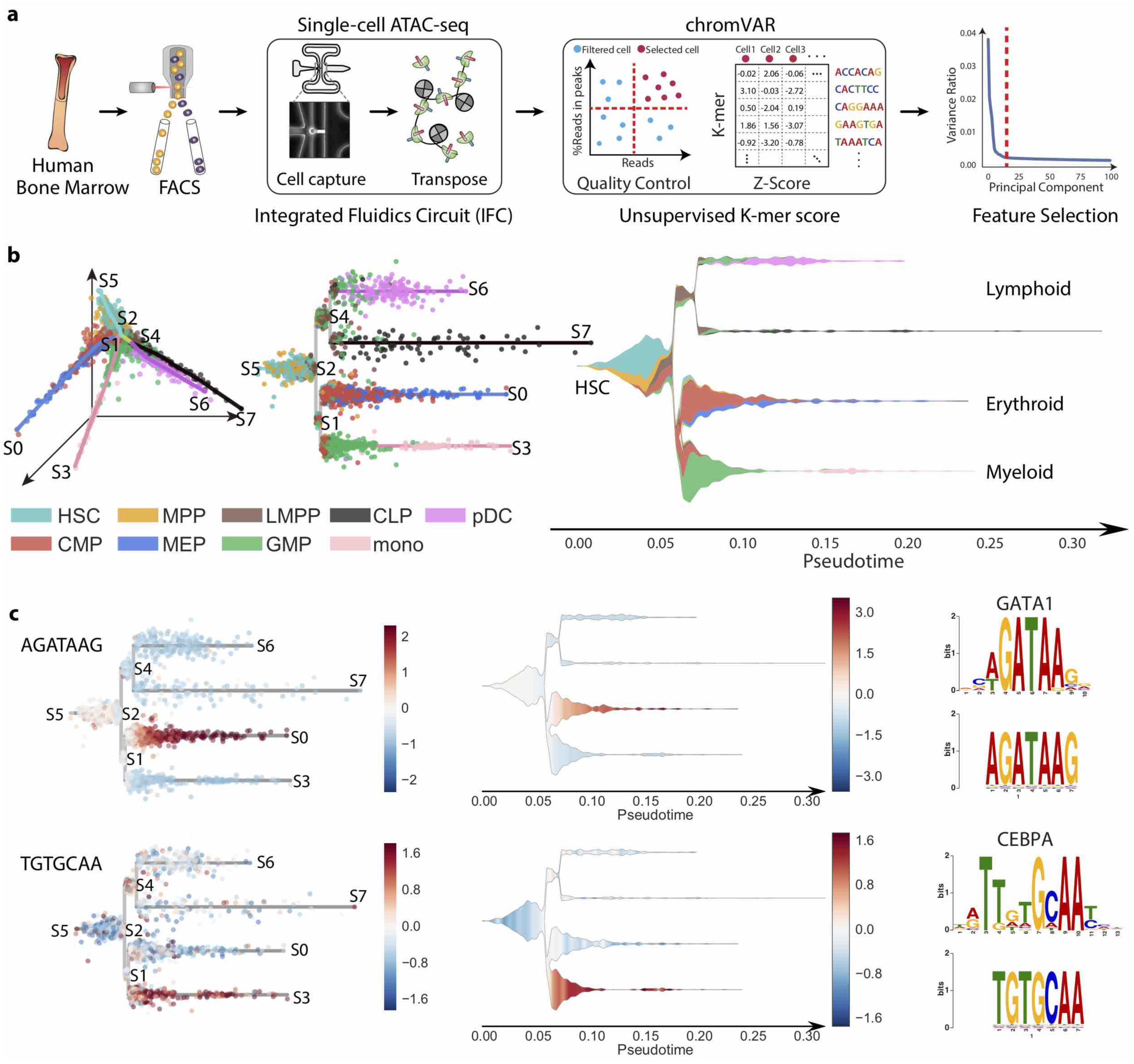
STREAM on single cell epigenomic data from the human hematopoietic system. **(a)** Single cell ATAC-seq workflow. FACS sorting is used to isolate populations from CD34+ human bone marrow and single-cell ATAC-seq measurements are performed. After mapping reads to the reference genome, reads within peaks are selected and ChromVAR is used to calculate k-mers’ z-scores. Finally, PCA is applied to the z-score matrix and top principal components are selected as features for STREAM analysis. **(b)** STREAM learns a principal graph from chromatin accessibility data and accurately reconstructs cellular developmental trajectories of the human hematopoiesis. As in Figure 1, the structure can be easily visualized thanks to the subway map and stream plots. In the first branch, the HSCs segregate through MPP into lymphocyte-committed, erythrocyte-committed and myelocyte-committed branches. STREAM also reconstructs the bifurcation from lymphoid multipotent progenitors (LMPP) to CLP and plasmacytoid dendritic cells (pDC). **(c)** Discovery of transcription factors important for lineage commitment. 7-mer DNA sequences are automatically detected and their frequencies are visualized in both the subway map and stream plots. Recovered 7-mer DNA sequences are mapped to known transcription factors motifs. We recovered *GATA1* and *CEBPA* as top hits, two classic master regulators in blood development, which correlate with directionality toward erythroid differentiation and myeloid differentiation, respectively.

To test the effectiveness of STREAM, we examined open chromatin profiles of > 2,000 cells profiled by scATAC-seq in known human hematopoietic lineages^20^. STREAM not only accurately reconstructs cellular developmental trajectories of the human blood system, but also recovers key sequence features and master regulators that have been implicated in differentiation and lineage commitment for different subpopulations **(Supplementary Fig. 15, Supplementary Note 6)**. For example, two of the detected 7-mer sequences match binding models for the transcription factors *GATA1* and *CEPBA*, which regulate differentiation towards erythroid and myeloid lineages, respectively **(Fig. 15b-c)**.

STREAM is available as user-friendly open source software and can be used interactively to explore several precomputed datasets and to compute new trajectories at stream.pinellolab.org **(Supplementary Fig 16**), or as a standalone command-line tool using Docker (github.com/pinellolab/stream) **(Supplementary Note 7-8)**.

## ONLINE METHODS

### STREAM framework

#### Trajectories inference

**Feature selection:** For transcriptomic data (single-cell RNA-seq or qPCR), the input of STREAM is a gene expression matrix, where rows represent genes, columns represent cells. Each entry contains an adjusted gene expression value (library size normalization and log2 transformation) **(Supplementary Note 7)**. The most variable genes are selected as features, using a procedure we have previously proposed^1^. Briefly, for each gene, its mean value and standard deviation are calculated across all the cells. Then a non-parametric local regression method (LOESS) is used to fit the relationship between mean and standard deviation values. Genes above the curve that diverge significantly are selected as variable genes.

**Dimensionality reduction:** Each cell can be thought as a vector in a multidimensional vector space in which each component is the expression level of a gene. Typically, even after feature selection, each cell has still hundreds of components, making it difficult to reliably assess similarity or distances between cells, a problem often referred as the *curse of dimensionality*^2^. To mitigate this problem, starting from the genes selected in the previous step we project cells to a lower dimensional space using a non-linear dimensionality reduction method called Modified Locally Linear Embedding (MLLE)^3^. MLLE takes into account local similarity of each cell with its neighbors and addresses the regularization problem of standard LLE by introducing multiple weight vectors in each neighborhood. The neighbor size is chosen based on the number of cells and is set by default to 10% of the total number of cells. The number of MLLE components to use depends on the number of branches and on the complexity of the structure to learn. Typically, three components capture the main structure for most datasets, increasing them may recover finer structures (although we observed that there is no benefit for selecting more than 5 components in all the datasets tested).

#### ElPiGraph: structure learning and fitting

##### Seeding initial tree structure

To create an initial seed structure for the principal graph learning by ElPiGraph we first used the affinity propagation^4^ method to cluster cells in the MLLE space. Affinity propagation is based on the idea of *message-passing* between sample points, and finds a small set of exemplars which are considered to be most representative of the other samples. For all our tests we used the scikit-learn implementation^5^ with a damping factor set to 0.75. Based on the exemplars obtained by the affinity propagation procedure, a minimum spanning tree (MST) was constructed using the Kruskal’s algorithm. The obtained tree is then used as initial tree structure for the ElPiGraph procedure.

##### Elastic principal graph method (ElPiGraph)

Elastic principal graphs are structured data approximators^6-8^, consisting of vertices and edges. The vertices are embedded into the space of the data, minimizing the mean squared distance (MSD) to the data points, similarly to *k*-means. Unlike unstructured *k*-means, the edges connecting the vertices are used to define an elastic energy term. The elastic energy term and MSD are used to create penalties for graph edge stretching and the bending of branches. To find the optimal graph structure, ElPiGraph uses a *graph grammar approach*, which is described below. This approach allows an effective exploration of the graph structure space via a gradient descent-like search. In STREAM, the set of graph grammars ^9^used always result in the construction of a principal tree (i.e., a graph without cycles) but alternative graph grammars can produce more complex (e.g., circular) topologies.

Let *G* be a simple undirected graph with a set of vertices *V* and a set of edges *E* and ϕ:*V* → **R**^*m*^ a map that describes an embedding of the graph into the multidimensional space **R**^*m*^ by mapping a node of the graph to a point in the data space. Let a *k*-star be a subgraph of *G* with *k* + 1 vertices *v*_0,1…,*k*_ ∈ *V* and *K* edges over these vertices {(*v*_0_, *v_i_*)|**i** = 1, ..,*k*}. Let *E*^(*i*)^(0), *E*^(*i*)^(l) denote two ends of the graph edge *E*^(*i*)^
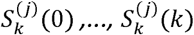
denote the vertices of a *k*-star
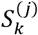
(where
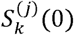
is the central vertex, to which all other vertices are connected). Let deg(*v_i_*) denote a function returning the order *k* of the star with the central vertex *v_i_* and zero if there is no any star centered in *v_i_*.

The *elastic energy of the graph embedment* is defined as the sum of squared edge lengths (weighted by the *λ_i_*, elasticity moduli and a penalty for excessive branching *α*) and the sum of squared *deviations from harmonicity* for each star (weighted by the *μ_j_*)

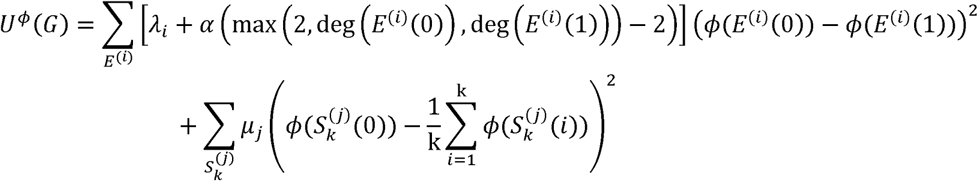

The second term (the deviation from star harmonicity) in the case of 2-star is a simple surrogate for minimizing the local curvature. In the case of *k*-stars with *k*>2 it can be considered as a generalization of local curvature defined for a branching point^8,10^.

Let *K* be a partition of all the data point under consideration (*X*_1_,*X*_2_,… *X_N_*) such that *K*(*i*) = argmm_*j*=1 … *N*_(*X_i_* − ϕ(*V_j_*))^2^ returns an index of the vertex in the graph which is the closest to the *i*th data point among all graph vertices. The objective function that we want to minimize is defined as

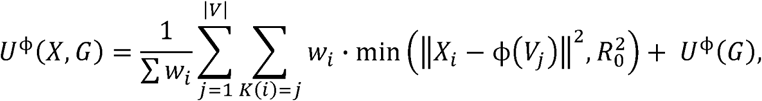

where *w_i_* is a weight of the data point *i* (can be unity for all points), |*V*| is the number of vertices, ‖..‖ is the usual Euclidean distance and *R*_0_ is a trimming radius that can be used to limit the effect of *points distant from the graph* (and hence to enforce a *local construction* that is more robust to noise) ^11^.

Given a graph topology for approximating a set of vectors *X*, our goal is to find a map ϕ:*V*→ **R**^*m*^ such that *U*^ϕ^(*X*,*G*) → min over all possible elastic graph *G* embedment in **R**^*m*^. The local minimum of *U*^ϕ^(*X*,*G*) is found by applying the usual splitting type algorithm:

1. Given the partition *K* of the data points by proximity to the graph vertices, we minimize *U*^ϕ^(*X*, *G*). Note that this functional is quadratic if *K* is fixed, therefore, the solution can be found very fast by solving a system of |*V*| linear equations.
2. Update *K* using new vertex positions. This simple step can be also implemented very fast.
3. Repeat 1) and 2) until a convergence criterion is met (i.e., the vertices are being displaced by less than a fixed threshold). Note that convergence is guaranteed by the form of *U*^ϕ^(*X*, *G*), which is a Lyapunov function wrt to the iterations 1-2.

A graph grammar-based approach for simultaneous learning of the graph topology and embedment of the graph into the data space starts from a seed graph *G*_0_and a map ϕ_0_(*G*_0_). A set of grammar operations are then applied iteratively to transform the graph topology, and hence the map, starting from a given pair {*G_i_*, ϕ*_i_*,(*G_i_*)}^12^. Each grammar operation *Ψ^p^* produces a set of s new candidate graph topologies *Ψ^k^*, possibly taking into account the dataset *X*:

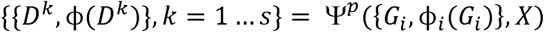

Given the pair {*G_i_*, ϕ*_i_*,(*G_i_*)} characterizing the *i^th^* step of the algorithm, a set of *r* different graph operations {Ψ^1^,…Ψ′}(which we call a “graph grammar”), and an energy function *U*^ϕ^(*X*, *G*), the algorithm applies all the grammar operations selected, fit the newly derived graph topologies to the data, and choose the most energetically favorable embedment as principal graph of the step (*i+l)^th^*:

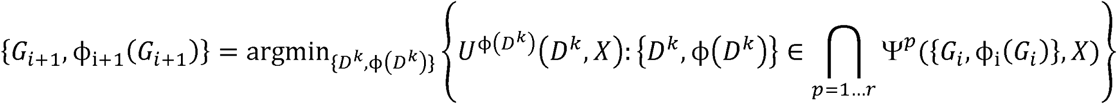

where {*D^k^*, ϕ(*D^k^*)} is supposed to be fit to the data after the application of a graph grammar.

In order to produce principal trees, one defines two operations for graph growth (‘bisect an edge’ and ‘add a node’) and one operation for graph shrinking (‘remove an edge’). Afterwards, two applications of growth operations are followed by one of shrinking. Such an approach allows avoiding local minima in the structure space of all possible tree topologies.

The ElPiGraph algorithm has four parameters with clear meaning and effect on the final result:

1. *λ* = *λ_i_*, controls the total length of the graph and, at the same time, promotes equal distance between neighbour graph nodes in the data space
2. *μ* = *μ_i_* controls the smoothness of the graph embedment (for the tree, smoothness of tree branches and harmonicity of graph stars).
3. *α* controls for excessive branching such that sufficiently large *α* (e.g., *α* = 1) hardly penalizes any branching while smaller values (e.g., *α* = 0.01) leads to keeping essential branches.
4. *R_0_* is the trimming radius, allowing robust estimation of node positions. ElPiGraph implements a simple scaling statistics allowing to automatically estimate *R_0_* if needed.

In the simplest case, *R_0_* = ∞, *α*= 0, and it is recommended to keep λ ≈ μ,/10. For all the single cell datasets in this paper it is desirable to set *α* = 0.02 and sometimes use trimming (automatically defined finite value for *R_0_*).

##### Adjusting the final tree structure

The resulting principal graph is refined based on the following procedures: 1) Principal tree branches can be extrapolated from the terminal vertices, i.e. a branch can grow, if necessary, to better fit cells that maybe fall far away from a terminal node. This allows a smoother pseudotime mapping and a more reliable characterization of cells close to initial or terminal points 2) Branches not supported by at least *n_minload_* data points can be removed or shrunk. 3) A *k*-star node, (node with connectivity *k*>2) can be rewired to another graph node if the latter has a higher local density or a larger number of cells projected into it to improve the positioning of candidate branching points.

In this paper, the *ElPiGraph.R* R package has been used, available at https://github.com/sysbio-curie/ElPiGraph.R. Implementations of the ElPiGraph are also available in other programming languages (Matlab, Java, Python, Sca1a)^13^.

#### Visualization

**Flat Tree Plot:** The tree structure learned in the 3D space (or higher dimensional space), is first approximated by linear segments (each representing a branch) and mapped to a 2D plane based on a modified version of the force-directed layout Fruchterman-Reingold algorithm^14^. In particular, we adjust each edge length in order to preserve the lengths of the branches of the original tree. Finally, using both the pseudotime location on the assigned branch and the distance from it in the MLLE space, we map cells to the obtained tree in the 2D plane. Cells are represented as dots and randomly placed to either side of the assigned branches. Each node in the tree indicates one cell state (cell states are sequentially named S0, S1, … starting from a randomly selected node) and the resulting structure is called *flat tree plot.*

**Subway map plot:** Starting from the flat tree plot and with a designated root or start node, breadth-first search is used to order and arrange nodes and edges horizontally on a 2d plane. Because we preserve the branch lengths of the original tree, the x-axis represents the distance (namely pseudotime) from the start node along the different branches. Cells are then mapped to the obtained structure, called *subway map plot* with the same strategy used for the flat tree plot. To display gene expression, each cell is colored according to its gene expression (the maximum value in the colormap is set as 90 percentile of gene expression values across all cells).

**Stream plot:** Starting from the subway map plot, for each cell type (if cell labels are provided), using a sliding window approach, we first calculate the number of cells in each window along a developmental branch. To provide smooth transitions around the branching nodes, in those regions the sliding window spans both parent branch and children branches and then proceeds independently on each branch. Then, the numbers of cells in all sliding windows are normalized based on the length of the longest path in the tree. The vertical layout of different branches is optimized by taking into consideration normalized numbers of cells to make sure there will not be overlap between branches. Based on the normalized sliding window values, we first use linear interpolation to construct a set of supporting points. Then the Savitzky-Golay filter (a smoothing filter able to preserve well the signal and avoid oscillations)^15^ is applied to create smooth curves based on the set of supporting points. Finally, the obtained curves polygons (one for each cell type) are assembled to form the *stream plot.* On stream plot, the length of each branch is the same as in the subway map plot and represents pseudotime, whereas the width is proportional to the number of cells at a given position. To display gene expression, we consider, for each sliding window, not only the number of cells but also their average gene expression values smoothed by bicubic interpolation (the maximum value is set as the 90th percentile of the average gene expression values from all the sliding windows).

#### Discovery of marker genes

**Diverging gene detection:** For each pair of branches *B_i_* and *B_j_*, and for the gene *E*, the gene expression values across cells from both branches are scaled to the range [0,1]. For gene expression *E_i_* from *B_i_* and gene expression *E_j_* from *B_j_*, we first calculate their mean values. Then, we check the difference between mean values to make sure it is above a specified threshold (the default value is 0.2). Mann-Whitney U test is then used to test whether *E_i_* is greater than *E_j_* or *E_i_* is less than *E_j_.* Since the statistic *U* could be approximated by a normal distribution for large samples, and *U* depends on specific datasets, we standardize *U* to Z-score to make it comparable between different datasets. For small samples where this test is underpowered (<20 cells per branch), we report only the fold change to qualitatively evaluate the differences between *E_i_* and *E_j_.* Genes with Z-score or fold change greater than the specified threshold (2.0 by default) are considered as differentially expressed genes between branches. Formally:

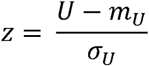

Where *m_u_*, *σ_U_* are the mean and standard deviation, and

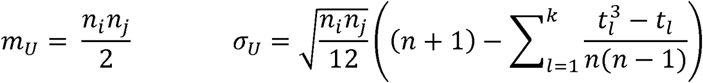

Where *n* = *n_i_* + *n_j_ n_i_*, *n_j_* are the number of cells in each branch, *t_i_* is the number of cells sharing rank *ℓ* and *k* is the number of distinct ranks.

**Transition gene detection:** For each branch *B_i_* and for each gene *E* we first scale the gene expression values to [0,1] for convenience. Then we check if the candidate gene has a reasonable dynamic range considering cells close to the start and end points. To this end, we consider the difference in average gene expressions of the first 20% and the last 80% of the cells based on the inferred pseudotime. If the difference is greater than a specified threshold (the default value is 0. 2), we then calculate Spearman’s rank correlation between inferred pseudotime and gene expression of all the cells along *B_i_.* Genes with Spearman’s correlation coefficient above a specified threshold (0.4 by default) are identified and reported as transition genes.

**Mapping procedure:** For a set of unmapped cells X = {*x_i_* | *i* = 1,…, *M*} and a fitted tree *T* built using the set of cells Y = {*y_j_*|*j* = 1,…,*N*} currently we have the assumption that X and Y have the same measured genes and are sequenced using the same experiment protocol. Both are X and Y are library size normalized and log2 transformed. To map cell *x_i_* into the embedding, we first find its nearest *K* neighbors in *Y*, based on the same feature genes and *K* used to build *T.* The largest distance between *x_i_* and its *K* neighbors is then chosen as the radius *r.* Then all the cells in *Y* within the radius*j_i_* = {*y_j_*| *d*(*x_j_*,*y_j_*) ≤ *r*) are used to compute a set of weights *W_i_* = {*w*_*ji*_, *j* ∈*j_i_*} as defined in the original MLLE procedure. Finally, using the MLLE embedding vectors *V* = {*v*_1_,…, *v_N_*}, the new cell position *x*′_*i*_ is calculated in the embedding with the following equation:

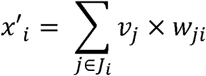

After mapping, each cell is assigned to its closest branch in T.

**STREAM analysis on scATAC-seq data:** For the scATAC-seq analysis, a total of 3,072 cells were profiled using FACS to isolate 9 distinct populations from CD34+ human bone marrow, which encompassed progenitors for four well-defined lineages^16^. 2,034 high-quality cells passed quality control filtering and were used in the downstream analysis with STREAM. Specifically, cells were filtered so that 1000 unique nuclear fragments were observed for each cell and at least 60% of these reads aligned in open chromatin peaks. After filtering low quality cells, the mean intensity and GC content for each peak that was called for this dataset was computed using the addGCBias function for the hgl9 genome using the BSgenome.Hsapiens.UCSC.hgl9 package available through chromVAR ^17^. These two coordinates were used to infer an empirically-defined set of background peaks to compute accessibility deviations, which have been described elsewhere^16,18^. As features we used an unbiased k-mer scoring, which is naive to any known transcription factor motif and thus generalizadle to other systems. We used the matchKmers function in chromVAR with parameters k = 7 and genome = BSgenome.Hsapiens.UCSC.hgl9, which returns a matrix of dimension number of peaks by number of k-mers where a 1 indicates that the peak contains the k-mer sequence. The output of this function was then included in the computeDeviations function to compute chromatin accessibility z-scores for each of the k-mers in our dataset. This matrix of cells by k-mer accessibility z-scores serves as a data-driven dimensionality reduction of the chromatin accessibility profiles of these cells. Based on the z-score matrix of k-mer DNA sequences, all the 7-mer features are standardized to have zero mean and unit variance. PCA is performed on the scaled matrix to convert z-score to principal components. According to the variance ratio elbow plot we selected the top 15 PCs, but excluded the first component since it captured technical noise (dropout and number of reads). The obtained matrix is used to reconstruct trajectories as previously described. Diverging and transition k-mers were selected with the same procedures used for gene selection. Finally, detected k-mers were mapped to known transcription factors using Tomtom ^19^(http://memesuite.org/tools/tomtom) and a motif database previously assembled [chromvar_and_hocomoco.meme](https://github.com/buenrostrolab/chromVARmotifs)^16^.

#### Comparison of methods for trajectory inference

**Simulated datasets:** Given a set of *n* cells and assuming we know their developmental/sampling time and topological organization, i.e. how they are organized in branches, we can easily evaluate a generic reconstruction method with the following two metrics:

1. Difference between the number of inferred and true branches.
2. Correlation between the true sampling time *X* and the inferred pseudotime *Y.* For the pseudotime we use either the proposed ranking or the actual distance from the starting point as provided by each method. We used 3 different measure of correlation: Pearson correlation *r*, Spearman correlation *ρ* and Kendall’s tau correlation *τ*, calculated as follow:

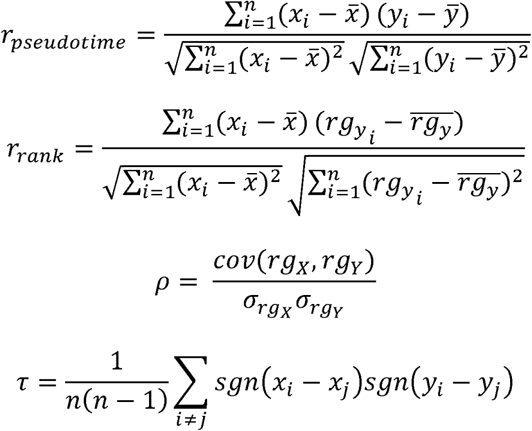

Where *rg_X_* and *rg_Y_* are the ranks of cells, *cov*(*rg_X_*,*rg*_Y_) is the covariance of rank variables, *σ*_rgx_and *σ_rgY_* are the standard deviations of rank variables. Note that since both Spearman correlation *ρ* and Kendall’s tau correlation *τ* are rank-based methods, the correlation between *X* and *Y* and the correlation between *X* and *rg_Y_* are the same, so we consider only the correlation between *X* and *Y.*

**Real datasets:** To evaluate the quality of reconstruction in real datasets in which we do not have the real developmental time and topological information, we used the following two metrics:

1. *Path-specific marker gene correlation analysis:* In real datasets oftentimes, we don’t have the sampling time along a branch. In this case, instead, it is helpful to evaluate how the inferred pseudotime recapitulates the progressive activation or repression of an important gene along that branch. The main idea here is that ordering cells based on a marker gene, which is important in defining a developmental trajectory, as a reasonable surrogate for the correct pseudotime ordering. As in the simulation case we computed 4 correlation coefficients using marker gene expression *X* and the inferred pseudotime *Y.*
2. *F*_1_ *score analysis on diverging or mutually exclusive marker genes:* Let us consider a pair of diverging or mutually exclusive marker genes, *G_i_* and *G_j_.* These genes should be highly expressed on different committed branches and rarely co-expressed in the same cell. We define *B_i_* as the branch which contains the most cells express *G_i_.* Then we can define as true positive (TP) for the number of cells expressing *G_i_.* The number of cells expressing *G_i_* on the other branches is defined as false negative (FN). The number of cells expressing *G_j_* on *B_j_* is defined as false positive (FP). Similarly, for *G_j_*, *B_j_* is the branch which has the most cells expressing *G_j_.* TP is the number of cells expressing *G_j_* on *B_j_.* FN is the number of cells expressing *G_j_* on the other branches. FP is the number of cells expressing *G_i_* on *B_j_.* Based on the following equations, recall, precision and FI score are calculated respectively for *G_i_* and *G_j_* as follow:

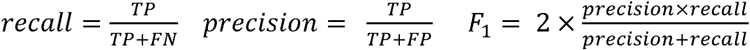

#### Website and code availability

STREAM is available as a user-friendly open source software and can be used interactively as a web-application at http://stream.pinellolab.org or as a standalone command-line tool: https://github.com/pinellolab/STREAM.

#### Data availability

All the data used in this study have been deposited at https://github.com/pinellolab/STREAM or available as
supplementary information.

## Supplementary materials

### Supplementary Note 1: STREAM analysis on scRNA-seq from the mouse hematopoietic system

We used STREAM to reanalyze scRNA-seq data from Nestorowa et al. 2016^1^, which sorted and profiled 1656 single cells including hematopoietic stem cells (HSCs), multipotent progenitors (MPPs), lymphoid multipotent progenitors (LMPPs), common myeloid progenitors (CMPs), granulocyte-monocyte progenitors (GMPs) and megakaryocyte-erythrocyte progenitors (MEPs), to study the mouse hematopoietic stem and progenitor cell differentiation processes.

STREAM recovers two bifurcation events (**Fig. 1b**) and three trajectories leading to myeloid, erythroid and lymphoid precursors. First, to check the validity of the structure, we used the assigned labels in the original study, which were derived by FACS sorting the different populations (when multiple labels were assigned based on different gates, we gave priority to narrow gates to obtain a unique label for each cell). As expected, HSCs progress into MPPs and then bifurcate into LMPPs and CMPs. Then CMPs differentiate into MEPs and GMPs respectively, hence accurately recapitulating known bifurcation events.

Second, using STREAM we rediscovered known marker genes, including diverging genes between two branches and transition genes along each branch. On the CMPs bifurcation which leads to GMPs (S3,S4) and MEPs (S3,S5) populations, STREAM detects diverging genes including GMPs-specific genes like Prtn3, Mpo, Epx, and MEPs-specific genes like Car2, Gata1, Mfsd2b^2 3,4^ (**Fig.lc, Supplementary Fig. 1a**). Along the MEP-committed trajectory (S3, S4), STREAM also recovered genes whose expression significantly correlates (p<lE-4, Spearman correlation) with the pseudotime progression. We recovered genes previously described ^5-7 8^ that are progressively and precisely downregulated such as Tmsb4x, Corola or upregulated like Blvrb, Ces2g (**Fig. 1c, Supplementary Fig. 1b**). Along the lymphoid differentiation trajectory (S2,S1,S0), STREAM identified HSCs-specific genes like Mpl^9^,Tgm2^10^ (**Supplementary Fig-2**), whose expressions are repressed towards lymphoid differentiation, and LMPPs-specific genes like lghv1-81, Ccl3 (**Supplementary Fig.2**), whose expressions are activated during the differentiation as discussed in the original study^1^.

Taken together, these analyses validate the accuracy of trajectory reconstruction of STREAM in recapitulating key bifurcation events and regulators of early blood development differentiation at single-cell resolution.

**Supplementary Figure 1.**
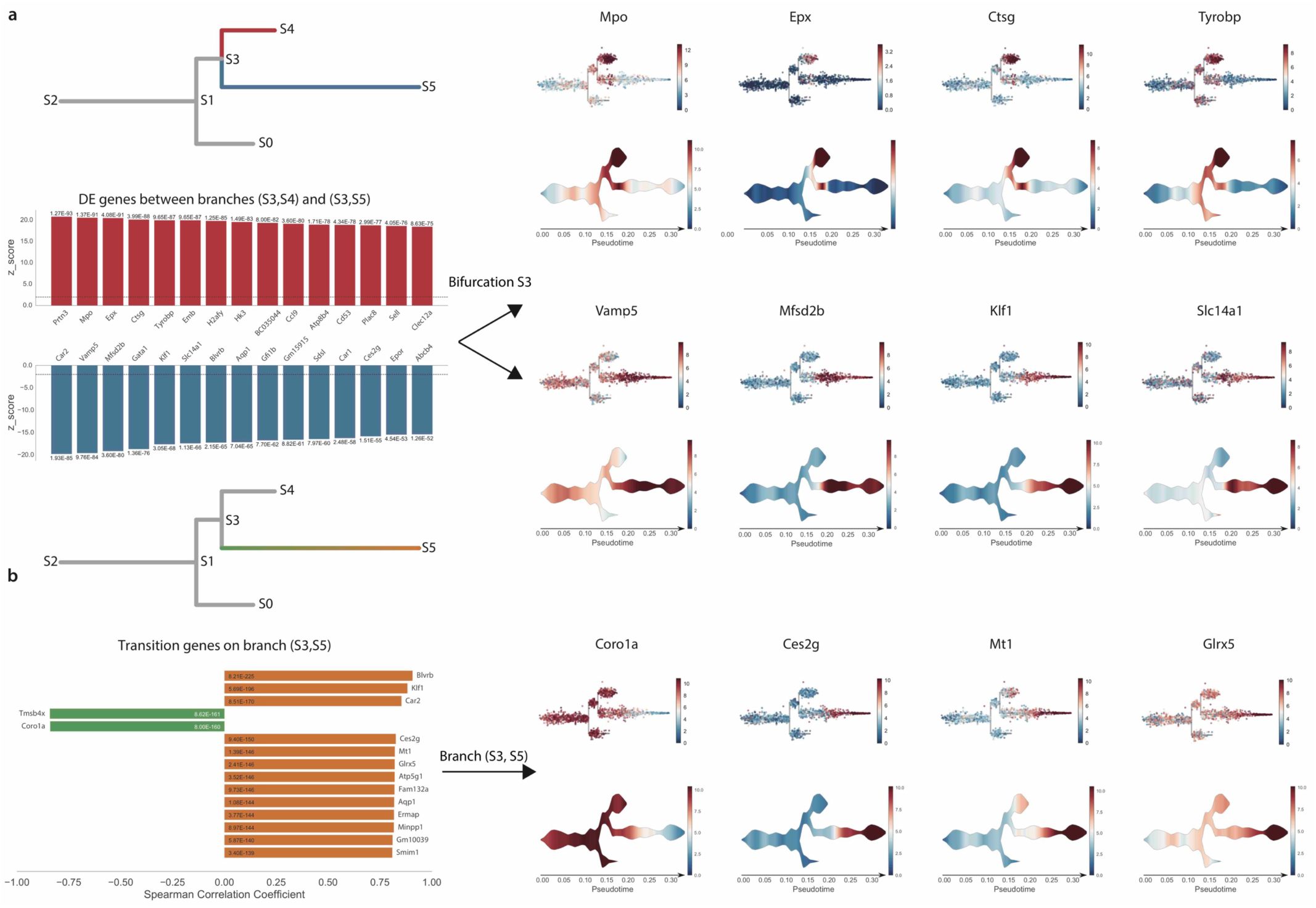
STREAM automatically discovers differentially expressed genes and transition genes around the SI bifurcation in mouse hematopoietic system. (a)Top left, subway map schematic to highlight the branches (S1,S3)(red), (S1,S0)(blue) used to calculate DE genes. Bottom left, genes highly expressed on branch (S1,S3) (red part) and genes highly expressed on branch (S1,S0) (blue part), sorted by significance. Top right, top detected marker genes for (S1,S3) are visualized on both subway map plots and stream plots. Bottom right, top detected marker genes for (S1,S0) are visualized on both subway map plots and stream plots. (b)Top left, subway map schematic used to highlights the branch (S1,S0) (green to orange gradient) used to calculate transition genes. Bottom left, genes monotonically increasing (orange part) or decreasing (green part) when progressing along branch (S1,S0), sorted by significance. Top Right, top detected upregulated and downregulated transition genes along branch (S1,S0) are visualized on both subway map and stream plots.

**Supplementary Figure 2.**
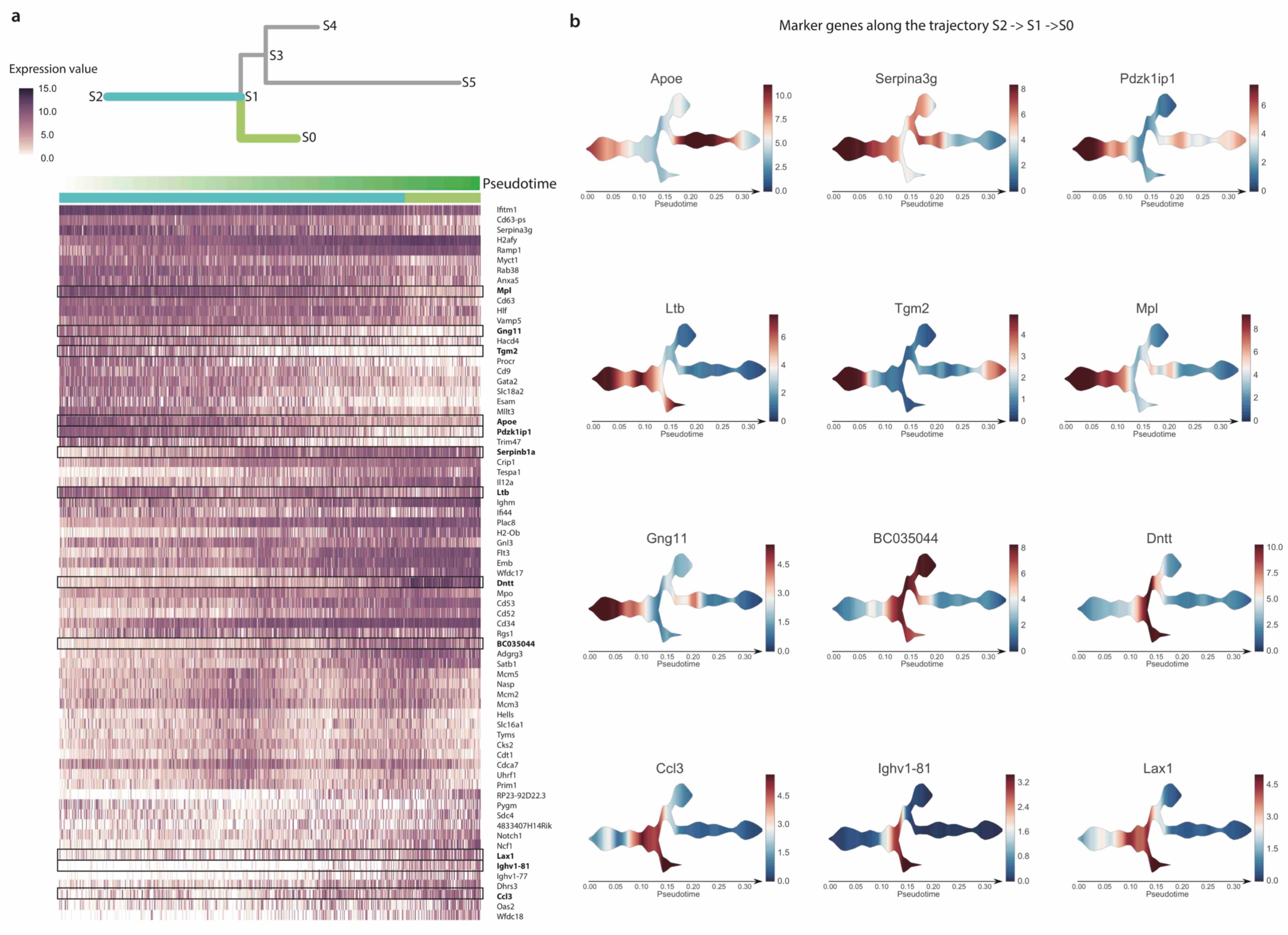
STREAM automatically discovers marker genes along lymphoid cells differentiation trajectory (S2,S1,S0) in mouse hematopoietic system. (a)Top left, subway map schematic to highlight the lymphoid cells differentiation trajectory (S2,S1,S0) consisting of two branches (S2,S1)(cyan), (S1,S0)(yellowgreen), whose related marker genes are calculated. Bottom left, heatmap showing the detected marker genes expression. Each row indicates one marker gene. Each column indicates one cell on the trajectory (S2,S1,S0). Cells are ordered by the inferred pseudotime from STREAM. The two colors represent cells’ branch ID assignment, either branch (S2,S1) or branch (S1,S0), (b) Twelve marker genes selected from left heatmap are visualized on stream plots.

### Supplementary Note 2: Studying genetic perturbations with STREAM’S mapping feature

STREAM provides a useful *mapping* procedure. After the principal graph is learned, it is possible to map new cells to the inferred structure. This *reference structure* is important when studying genetic or epigenetic perturbation, or when comparing different conditions (for example normal and cancer, response to stimuli, etc.). Existing methods require to fit a new model since the fitting procedure is not deterministic or because they don’t implement and provide this feature to the users. The main problem with re-computing the structure lies in the fact that it is hard to interpret pseudotime and cell positioning since trajectories may change based on the density and/or composition of the new cells to map. Our mapping procedure is instead deterministic and allows the user to easily study and predict perturbation effects, and explore the origin of unknown cell populations on annotated branching structures or vice versa (see an example in **Supplementary Note 3**).

To show the utility of the mapping feature, we applied STREAM on scRNA-seq data from Olsson et al. 2017^11^. Using FACS sorting, 382 cells were isolated and profiled from different subpopulations, including stem/multipotent progenitor (LSK; lin-, Scal+, c-Kit+), CMP, GMP, and LKCD34+(lin-c-Kit+CD34+) cells (**Supplementary Fig. 3a left**). A key result of this study is the discovery of metastable mixed-lineage states and the presence of co-expressed genes at single-cell level from competing lineages. The authors suggest that these metastable states are important in cell-fate decisions and that transcription factors play a key role in this process. In fact, they uncovered and validated two key transcription factors, i.e. Gfi1 and Irf8, that are co-expressed in a sub-population and are shown to be important for the commitment to neutrophils or macrophages. Importantly, this dataset contains, in addition to wild-type data, genetic perturbations of those two key regulators.

Using the wild-type data, STREAM unbiasedly and correctly reconstructs the cell lineage hierarchy (**Supplementary Fig. 3a** right) as shown by inspection of the labels proposed in the original study (either cell surface markers or predicted lineages): starting from haematopoietic stem cell/progenitor(HSCP), cells bifurcate into an erythrocytic branch (which contains megakaryocytic (Meg) and erythrocytic (Eryth) cells) and into a multi-lineage primed (Multi-Lin) branch. Multi-Lin cells further separate into the granulocytic (Gran) branch and monocytic (Mono) branch. The hierarchical progression can be easily visualized by our proposed 2D visualizations: subway map and stream plots (**Supplementary Fig.3b**). Importantly, STREAM precisely recovers the bifurcation event from Muli-lineage to Mono and Gran as shown in the original study within the wild-type GMP cellular population (**Supplementary Fig. 3b,c**), whereas the proposed Monocle2 analysis in the same dataset^12^ incorrectly assigns those cells to a very short erythroid branch. Furthermore, Monocle2 branch lengths are overall very diverse and distorted in their hierarchical representation (FE branch in Fig 2, and Supplementary Fig. 18 of the original paper^12^). Using our gene expression analysis, the Gran-specific gene Gfi1, Monospecific gene Irf8, and Eryth-specific gene Gata1 are highly expressed on their respective inferred trajectories, confirming the validity of the reconstructed branching structure (**Supplementary Fig. 3d**).

**Supplementary Figure 3.**
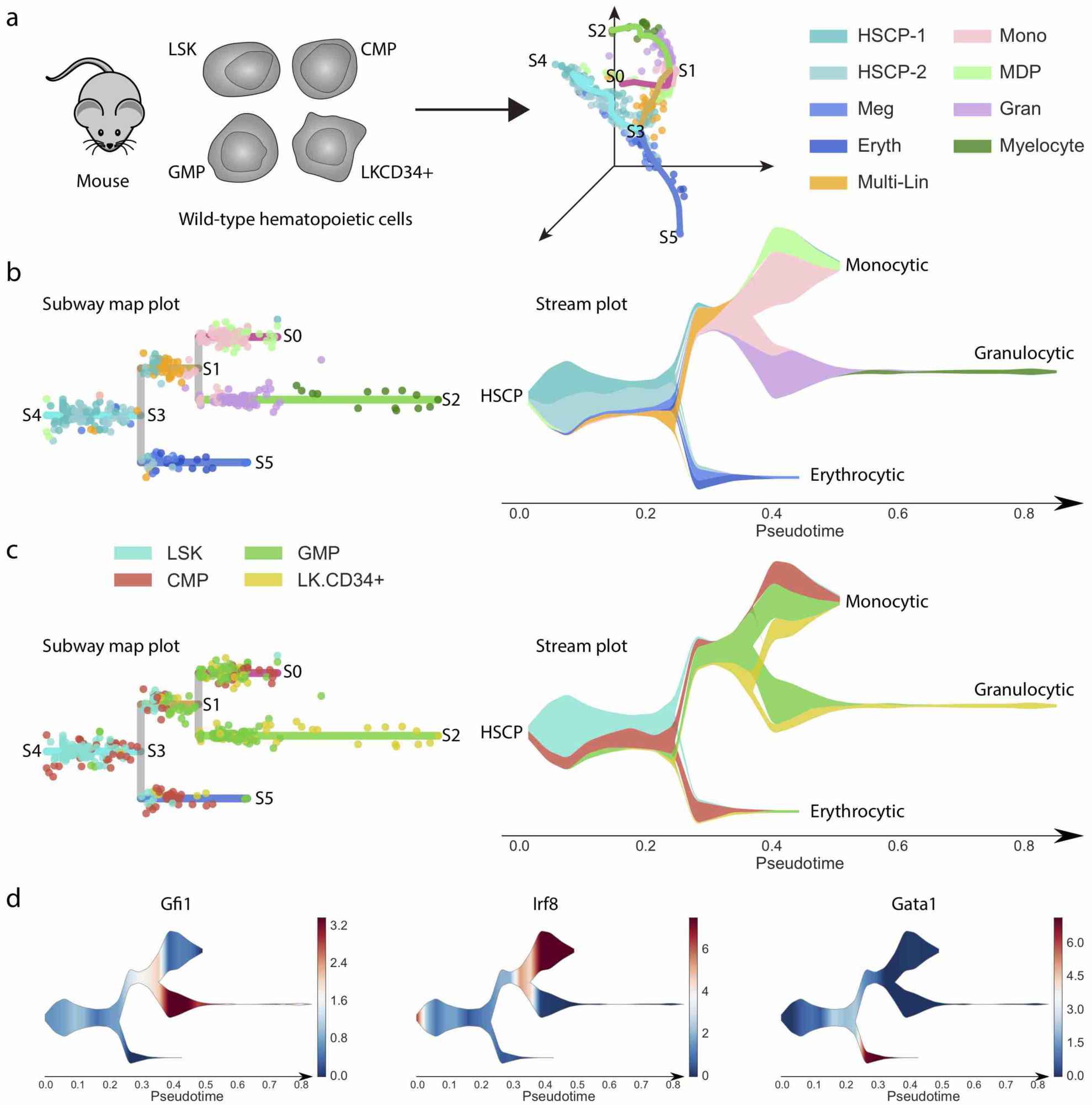
Unbiased reconstruction of wild-type mouse hematopoietic cell lineage hierarchy. (a) Left, cell subpopulations were isolated from the mouse wild-type hematopoietic system including stem/multipotent progenitor (LSK), common myeloid progenitor (CMP), granulocyte monocyte progenitor (GMP) and LKCD34+ and profiled by scRNA sequencing. Right, wild-type cells are shown in low dimensional space together with trajectories inferred by STREAM. Cells are colored by the cluster labels proposed by *Olsson at al*. (b) Left, subway map plot; right, stream plot. Both are colored by cluster labels inferred by *Olsson at al*. (c) The same subway map and stream plot as (b) but colored by FACS gating labels from the original study. (d) Stream plots of three key marker genes: Gfi1 for granulocyte, Irf8 for monocyte and Gata1 for Meg and Eryth.

Next, using the STREAM mapping function, we analyzed the genetic perturbation data to study the consequences on cell fate determination of Gfi1 loss (Gfi1 -/-), Irf8 loss (Irf8 -/-) and both Gfi1 and Irf8 loss (Gfi1 -/- Irf8 -/-) within wild-type GMP cells (**Supplementary Fig. 4a**). Gfi1 -/- GMP cells tend to differentiate into the Mono branch and Irf8 -/- GMP cells lean toward the Gran branch. Gfi1 -/- Irf8 -/- GMP cells have equal chance to go either way. The loss of Gfi1 and Irf8 instead does not show any imbalance of cells differentiating into the diverging branches (**Supplementary Fig. 4b-d**). Our predictions are validated by the original study where the authors used GMP cells with inducible expression and GFP reporters for Gfi1 and Irf8. Irf8 loss led to cells that differentiated toward granulocyte. Conversely, Gfi1 loss led the cells to differentiate toward monocytes. Interestingly they showed that cells from the hematopoietic stem cell/progenitor and myeloid compartments are trapped with the double knockouts of Irf8 and Gfi1, and in fact, are rarely differentiating towards monocytes or granulocytes. These results are in perfect agreement with our unbiased analysis. In addition, compared to the proposed Monocle2 analysis, our reference structure can be fixed to recapitulate only the wild-type cells and is not influenced by the mapping of new cells (compare instead A,B with C,D in Supplementary Fig. 18 of the original paper^12^).

**Supplementary Figure 4.**
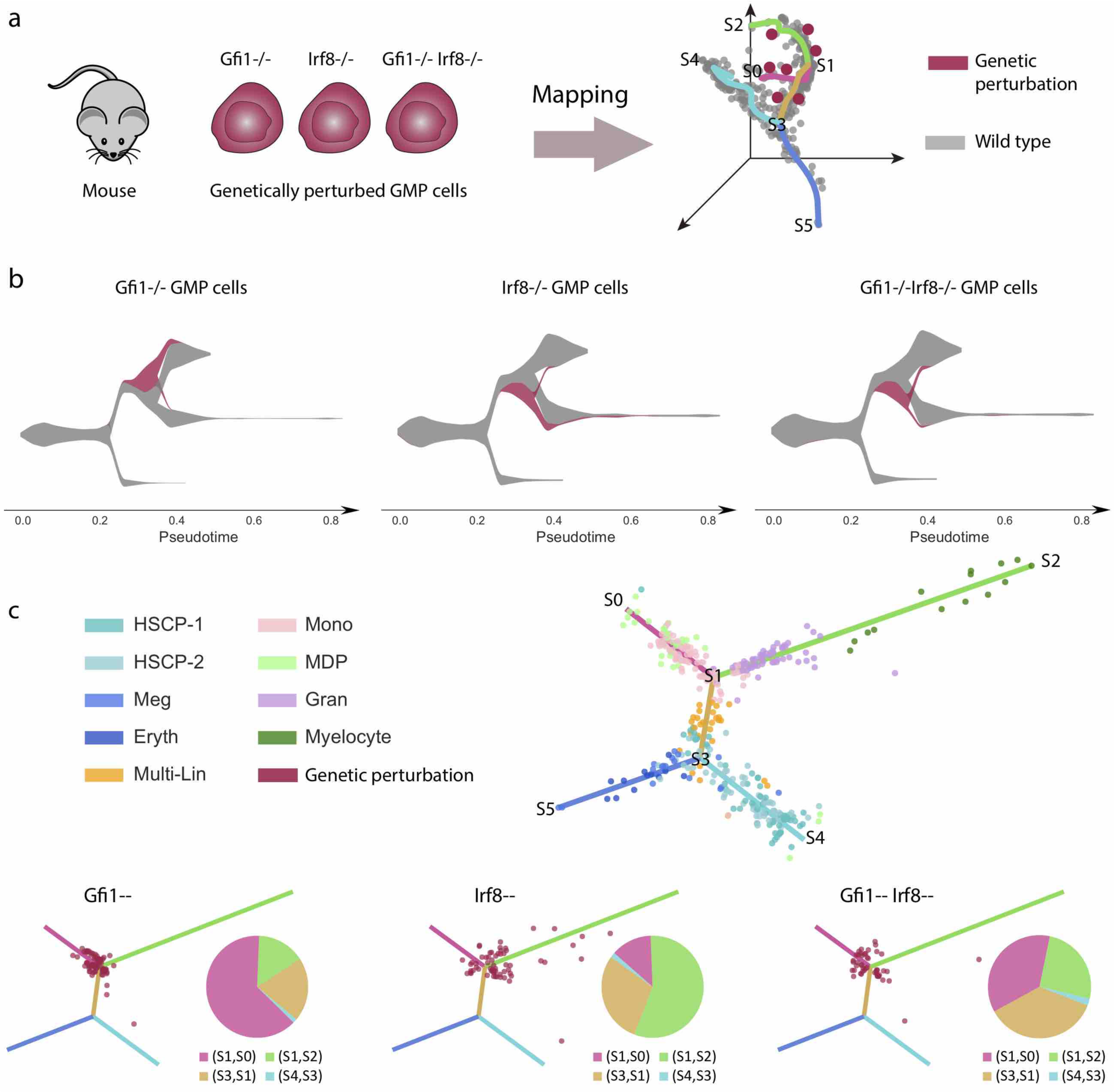
Mapping of genetic perturbation data to the inferred trajectories from wild-type mouse hematopoietic cells. (a) Left, scRNA sequencing is performed on genetically perturbed cells within the GMP populations: Gfi1-/-, Irf8-/- and Gfi1-/-Irf8-/-. Right, genetically perturbed cells are mapped using STREAM to the low dimensional space in which cellular trajectories were built based on wild-type cells, (b) At density level, stream plots easily summarize the effects of the 3 genetic perturbations: Gfi1-/-cells are diverted to monocyte-committed branch while Irf8-/-cells are instead diverted to granulocyte-committed branch. Gfi1-/-Irf8-/-cells have equal chances to differentiate into either branch, (c) Single-cell level visualization of perturbed cells on the reference flat tree plot constructed from wild-type cells (top). Genetically perturbed cells are mapped to the flat tree and shown in red. Pie charts show the proportion of genetically perturbed cells on different branches. Consistently with the stream plot in (b), Gfi1-/-cells mainly appear on monocyte-committed branch (S1,S0), while the majority of Irf8-/-cells appear on granulocyte-committed branch (S1,S2). Gfi1-/-Irf8-/-cells are approximately equally located on the intermediate state branch (S3,S1), monocyte-committed branch (S1,S0) and granulocyte-committed branch (S1,S2).

### Supplementary Note 3: STREAM analysis on single cell qPCR data of the zebrafish hematopoietic system

Next, we tested STREAM with data from a different organism, analyzing recently published single-cell qPCR data from Moore et al^13^, that provided a first model of the zebrafish hematopoiesis system using a panel of 96 gene primers. 166 cells were profiled from the wild-type(WT) whole-kidney marrow (WKM). STREAM analysis uncovered four cell lineages trajectories (**Supplementary Fig. 5a left**). Based on the automatic gene detection module of STREAM, we uncovered marker genes for each trajectory (**Supplementary Fig. 5a**), which includes T cell marker gene *TCR*-*alpha*, myeloid marker gene *nccrp*-*1*, B-cell marker gene *CD79* and erythroid marker gene *band3.* Based on this analysis, we hypothesize that the inferred four branches corresponded to T cells, myeloid, B cell and erythroid lineages (**Supplementary Fig. 5a** *right*).

**Supplementary Figure 5.**
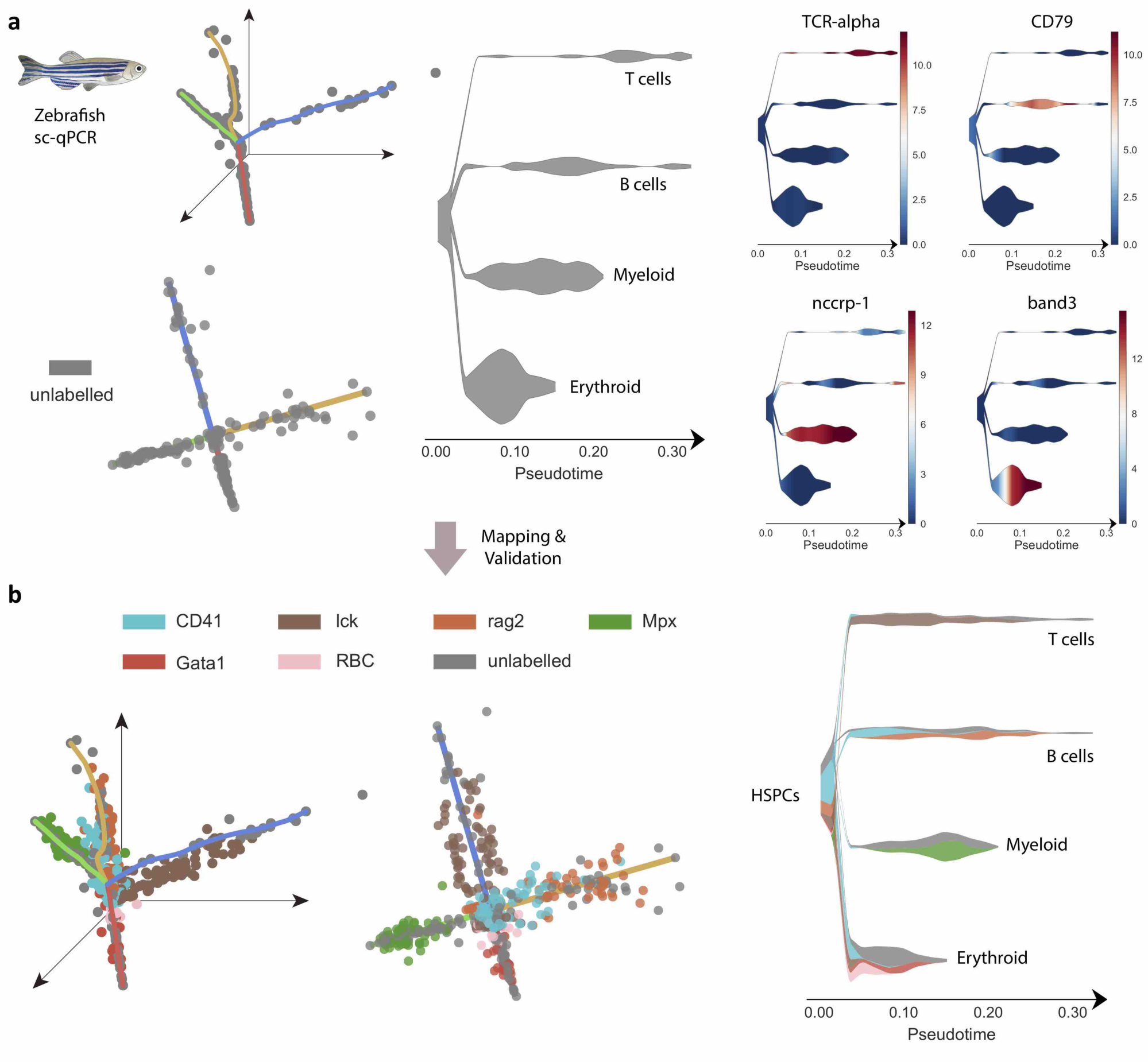
STREAM recovers developmental trajectories of hematopoietic cells in zebrafish from qPCR data. (a) STREAM output for single-cell qPCR on cells from zebrafish wild-type whole-kidney marrow (WKM). The stream plot shows only one color (gray) since no labels to annotate cell-types are available in this case. Four trajectories are recovered and visualized in the 3D space, flat tree plot and stream plot respectively. Four of the top marker genes automatically detected by STREAM are visualized as stream plots: TCR-alpha (T cells), cd79 (B cell), nccrp-1(myeloid) and band3(erythroid). (b) Validation of the putative cellular differentiation branches. Hematopoietic cells from adult transgenic zebrafish and peripheral blood are mapped to the trajectories inferred in (a). These cells comprise peripheral red blood cells (RBC) and FACS-sorted cells, which include CD41 (hematopoietic stem and precursor cells), lck (T cells), Mpx (myeloid cells), rag2(B cells) and Gata1 (erythroid cells).

To validate our branches, we used the STREAM mapping features to map fluorescent-labeled and FACS sorted cells from WKM: 20 erythroid cells from peripheral blood cells (per RBC), 24 erythroid cells Tg(gata1:dsRed), 48 myeloid cells Tg(mpx:GFP), 49 B cells Tg(rag2 :dsRed), 83 mature T cells Tg(lck:GFP)cells, 85 HSPCs Tg(CD41:GFP)low. Lck cells are mapped to T cell branch, mpx cells are mapped to myeloid cell branch, rag2 cells are mapped to B cell branch, both gata1 and per RBC are mapped to erythroid branch; also, the majority of HSPCs are mapped to the proposed starting state as expected (**Supplementary Fig.5b**). This analysis validates our unbiased reconstruction of the developmental trajectories from unlabeled cells and provides another example of the STREAM mapping feature utility.

### Supplementary Note 4: STREAM analysis on inDrop data of the zebrafish hematopoietic system

To test the scalability and robustness of STREAM on a larger and more challenging scRNA-seq dataset, we next analyzed “10000 unlabeled cells from the zebrafish whole-kidney marrow generated by Tang et al^14^ using the inDrop protocol^15^ (a recently proposed droplet-based assay). The original study, based on dimensionality reduction and clustering, uncovered and annotated 10 different and imbalanced subpopulations (validated by the authors using sorting of fluorescent or transgenic cell sub-populations) (**Supplementary Fig.6a**). STREAM correctly recapitulated the hierarchy of the different lineages and unbiasedly recovered four different hematopoietic cellular trajectories: starting from HSCs, through blood progenitor cells, cells differentiate into erythroid, macrophage, neutrophil and lymphoid lineages (**Supplementary Fig.6b**). Importantly, we rediscover well-known marker genes: *ba1* for the erythroid branch, *mfap4* for the macrophage branch, *mpx* for the neutrophil branch, and *lck* for the lymphoid branch (**Supplementary Fig.6c**). This analysis highlights four important points of our approach: 1) we can recover trajectories using unsorted populations, 2) the trajectory inference is robust to sub-populations imbalance, 3) our gene analysis is a powerful tool to discover marker genes, and 4) our method is scalable to currently available large-scale single-cell assays.

**Supplementary Figure 6.**
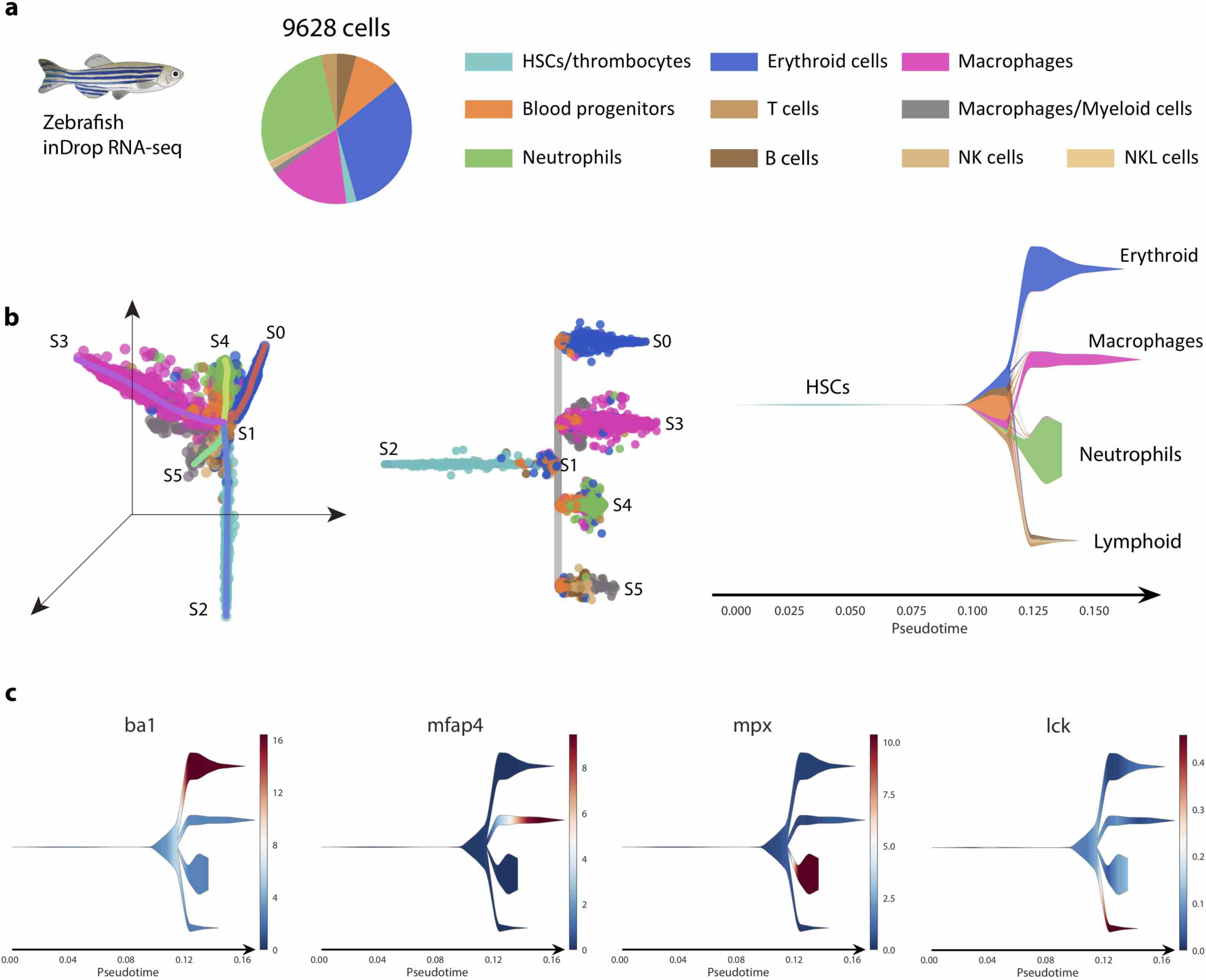
STREAM reconstructs cellular heterogeneity within the zebrafish kidney marrow from inDrop data. (a) STREAM output for inDrop single-cell RNA-seq data from the zebrafish wild-type whole-kidney marrow. Cell labels are based on the *Tang et at.* classification and are highly unbalanced as shown by the pie chart. (b) Principal graph plot, subway map plot and stream plot show the trajectories recovered in the hematopoiesis of zebrafish. HSCs through blood progenitor cells differentiate into erythroid, myeloid (including macrophage and neutrophil) and lymphoid cells (c) Marker genes from original study are visualized in stream plot to confirm and validate the recovered structure.

### Supplementary Note 5: Comparison of STREAM with existing methods

Several methods have been proposed for pseudotime inference or trajectory reconstructions. In fact, more than 50 methods have been proposed for this task making a systematic comparison unfeasible for the scope of this manuscript. For this reason, we compared STREAM with 10 state-of-the-art methods well recognized and commonly used by the single-cell community: Monocle2, scTDA, Wishbone, TSCAN, SLICER, DPT, GPFates, Mpath, SCUBA and PHATE^12,16-24^.

**Supplementary Figure 7.**
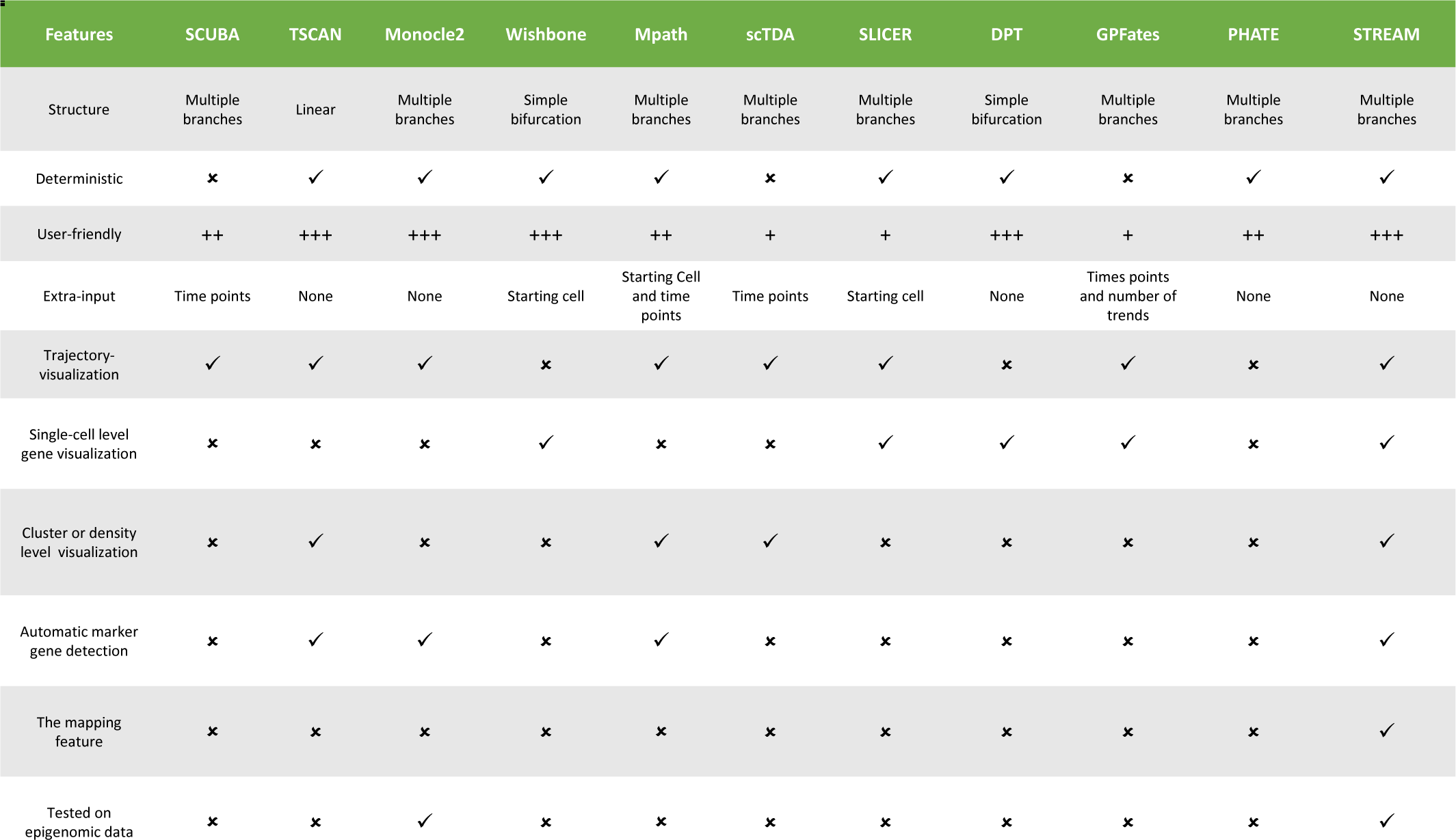
Overview of trajectories inference methods included in the comparison. Summary table to compare features available in different methods.

First, we reviewed the different methods to summarize their general features, required inputs, supported data and limitations (**Supplementary Fig. 7**). Next, we focus our comparison on three important aspects: topology correctness, pseudotime reliability, and branching model complexity. We also provide in our assessment the default visualizations provided by each method to showcase and easily compare their expressiveness in representing cellular development trajectories. For each method, the analyses were performed with standard parameters when possible, following the guidelines provided in the documentation or suggested by the respective authors.

We started our analysis using a previously proposed gold-standard synthetic dataset by Rizvi et al.^16^ with known topology and pseudotime: two bifurcation events and 3 different time points (**Supplementary Fig. 8**). We first assessed the qualitative output of each method using their proposed visualization. PHATE, a dimensionality reduction method, qualitatively preserves cellular trajectories but does not provide branch assignment information for cells. STREAM, scTDA, Monocle2, and Mpath can accurately reconstruct two bifurcation events. Wishbone and DPT instead can only detect one simple bifurcation even though the obtained 2D manifold clearly shows two bifurcations. SLICER detects too many branches and their positions are difficult to assess from the proposed visualization. GPFates requires to prespecify the number of trajectories (referred as trends in the original study and manually set for this dataset to 3). Although the generated curves initially follow the correct branches, they incorrectly converge at the end. SCUBA fails to detect bifurcation events in this dataset. We also noticed that scTDA, SCUBA, and MPATH do not provide single-cell-resolution visualization. Wishbone and DPT do not provide trajectory visualization so they cannot visualize both time points and recovered branch assignment in the same plot. (**Supplementary Fig.8**).

**Supplementary Figure 8.**
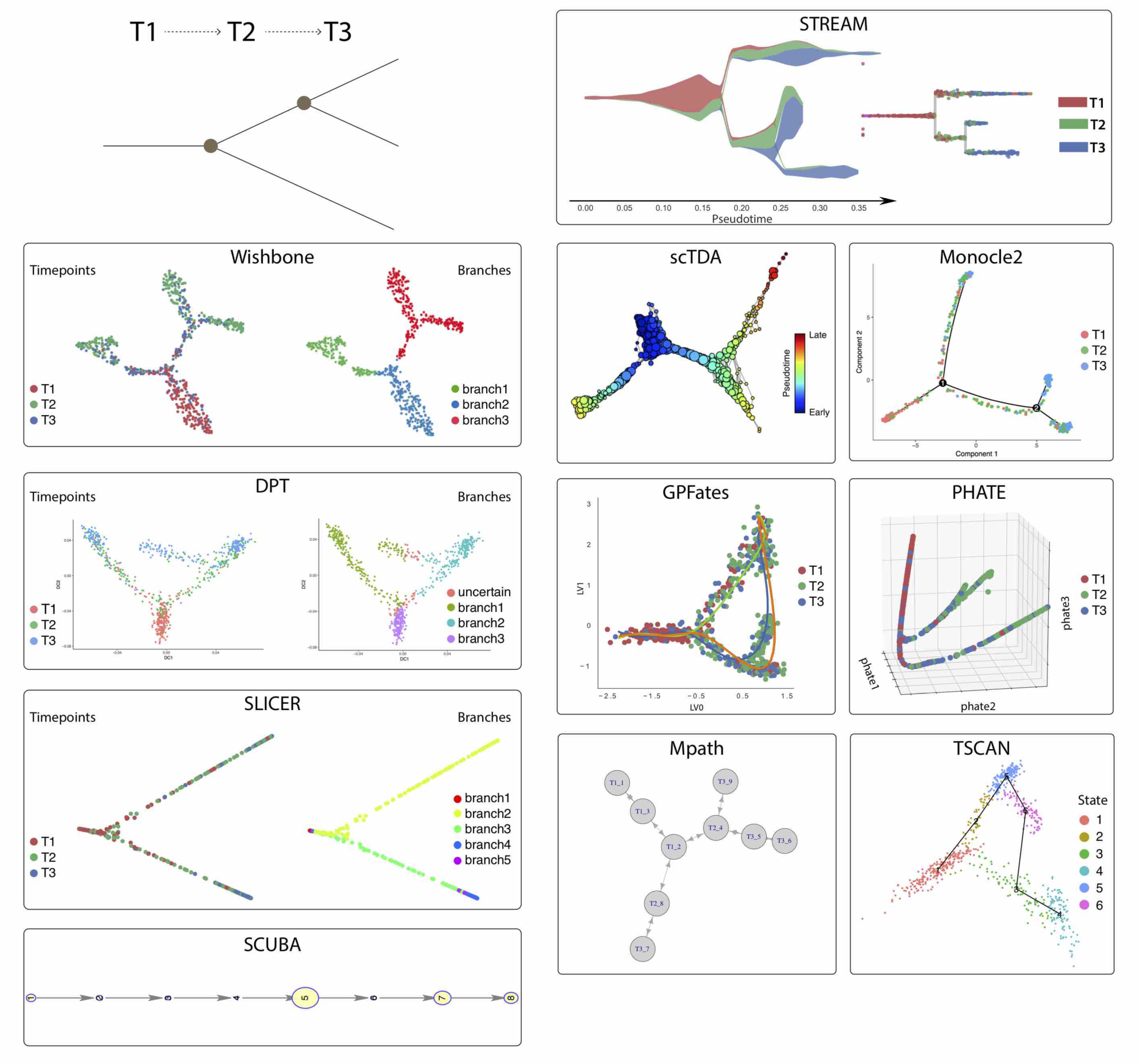
Output of different trajectories inference methods on synthetic data. Top left: Topology structure of a synthetic benchmark dataset proposed by *Rizvi et al*. Cells are sampled from three different time points (T1, T2 and T3). It has two bifurcation events: the first happens between T1 and T2, the second happens between T2 and T3. **STREAM**, stream plot and subway map plot allow to study cellular trajectories and time points at both density level and single cell level. **Wishbone**, left, cells are colored by time points, right, cells are colored by its identified differentiation branch ID. (inferred trajectories cannot be explicitly visualized). **scTDA**, proposed topological representation colored by pseudo-time indicates cellular differentiation trajectories, in which nodes correspond to a set of cells and node size is proportional to the number of cells (visualization of single-cells is not available). **Monocle2**, cells are colored by time points and the skeleton depicts differentiation trajectories (density information is not available). **DPT**, on the left cells are colored by time points and on the right cells are colored by its identified differentiation branch ID (inferred trajectories cannot be explicitly visualized). **GPFates**, cells are colored by time points. Three curves represent three trajectories(trends). (density information is not available). **PHATE**, cells are colored by time points. It doesn’t provide trajectories information and pseudotime. **SLICER**, on the left cells are colored by time points and on the right cells are colored by its identified differentiation branch ID (inferred trajectories cannot be explicitly visualized). **Mpath**, each node represents one landmark cell and the tree structure shows differentiation trajectories (visualization of single-cells is not available with this method). **TSCAN**, cells are colored by states detected from TSCAN. The skeleton depicts a linear trajectory. (density information is not available). **SCUBA**, each node is one cluster and the node size represents the variance within each cluster. (neither visualization of single-cells or density information is available with this method).

We next calculated for each method the correlation (**Online methods**) between true pseudotime and inferred pseudotime. Our method has the best performance for two out of four metrics (importantly when using the actual pseudotime defined as the distance of each cell from the origin in the proposed embedding) and comparable performance for the other two rank-based metrics (following scTDA in which this synthetic dataset was proposed) (**Supplementary Fig. 9**).

**Supplementary Figure 9.**
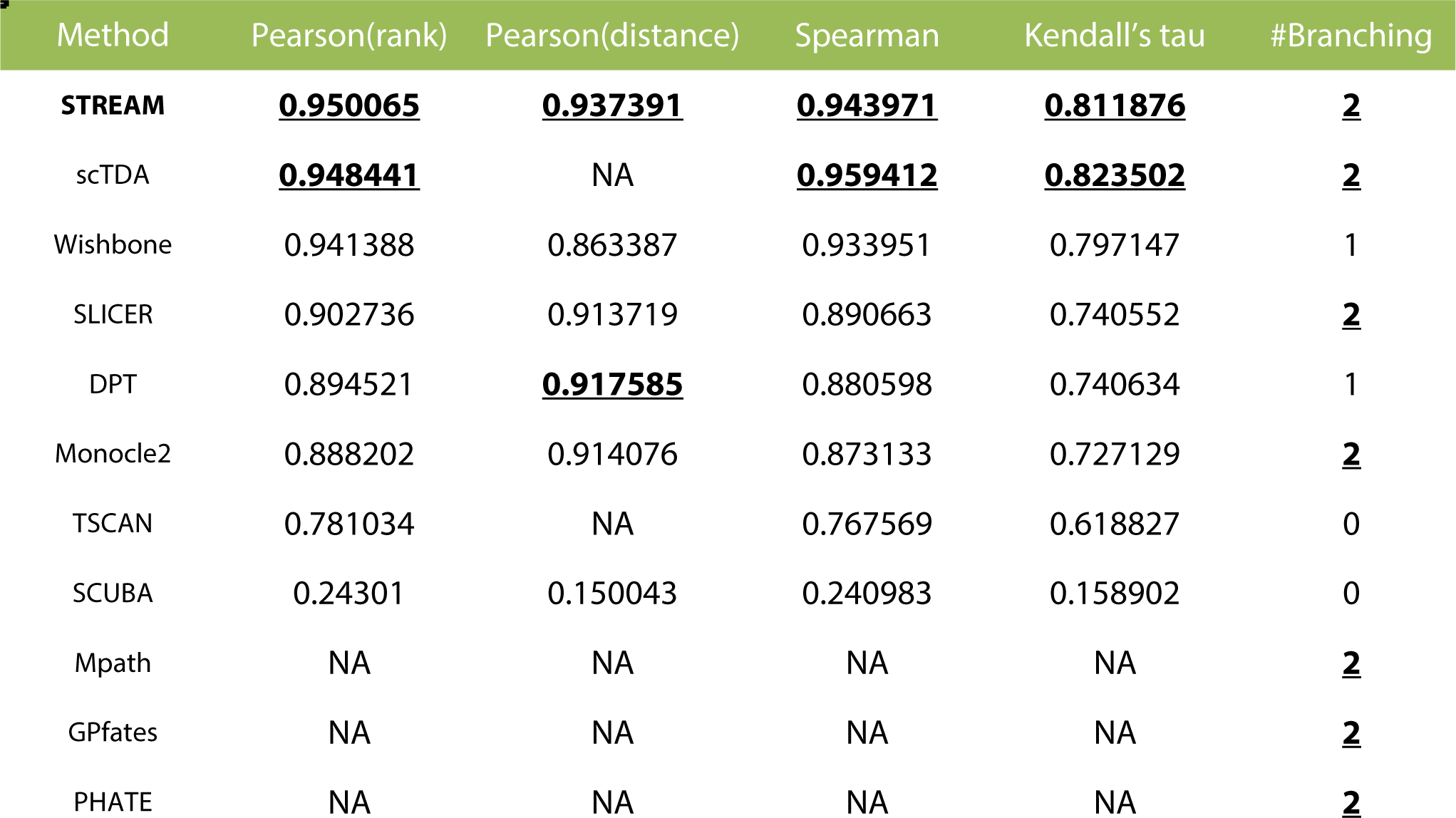
Analysis results of different methods on synthetic data. Correlation (Pearson, Spearman and Kendall’s tau) between real and inferred pseudotime (either rank or distance based) for the 11 methods tested.

In summary, STREAM correctly recovers the correct topology, has the best pseudotime reconstruction performance when using distance-based pseudotime along a trajectory, and is the only tool that provides a visualization to study the density of different cell types in different branches.

To compare the different methods on real datasets, we first used the most commonly used scRNA-seq dataset for this task, originally proposed by Trapnell et al^25^. This dataset contains human skeletal muscle myoblasts (HSMM) cells differentiating along a linear trajectory. In this analysis, we were able to evaluate only methods capable to detect the correct bifurcation event. Regardless, the visual output of all the methods is presented for completeness (**Supplementary Fig. 10**). The original study proposed a single bifurcation, which leads to myoblast cells or separate potentially contaminating cells (**Supplementary Fig. 11a**). To test the quality of pseudotime it has been proposed to use known marker genes along the myoblast differentiation trajectory, in particular to correlate their expression level (a surrogate for the correct ordering) with the rank or distance-based pseudotime (**Online Methods**). To this end, we used the previously proposed genes ENO3, MEF2C, and MYH2 ^12,18^. STREAM has the overall best performances on ENO3 and MYH2 (when considering the average score of the four metrics) and a comparable performance on MEF2C with both distance or rank based pseudotime (**Supplementary Fig. 11b-c, Supplementary Fig. 12**). When ordering cells based on the distance pseudotime, we expect, in the ideal scenario, a continuous and smooth distribution. For example, STREAM can generate a smooth and monotonically increasing distribution of ENO3 expression based on the inferred pseudotime. In contrast, we noticed that for the distance-based pseudotime in Monocle2, cells are mainly attracted to the end points of the trajectory, with few cells in between. In Wishbone and SLICER, distance-based pseudotime shows a set of unexpected discrete segments. Neither Mpath nor TSCAN can generate distance-based pseudotime. In addition, Mpath doesn’t recover a monotonically increasing trend (**Supplementary Fig. 11c, Supplementary Fig. 12**).

**Supplementary Figure 10.**
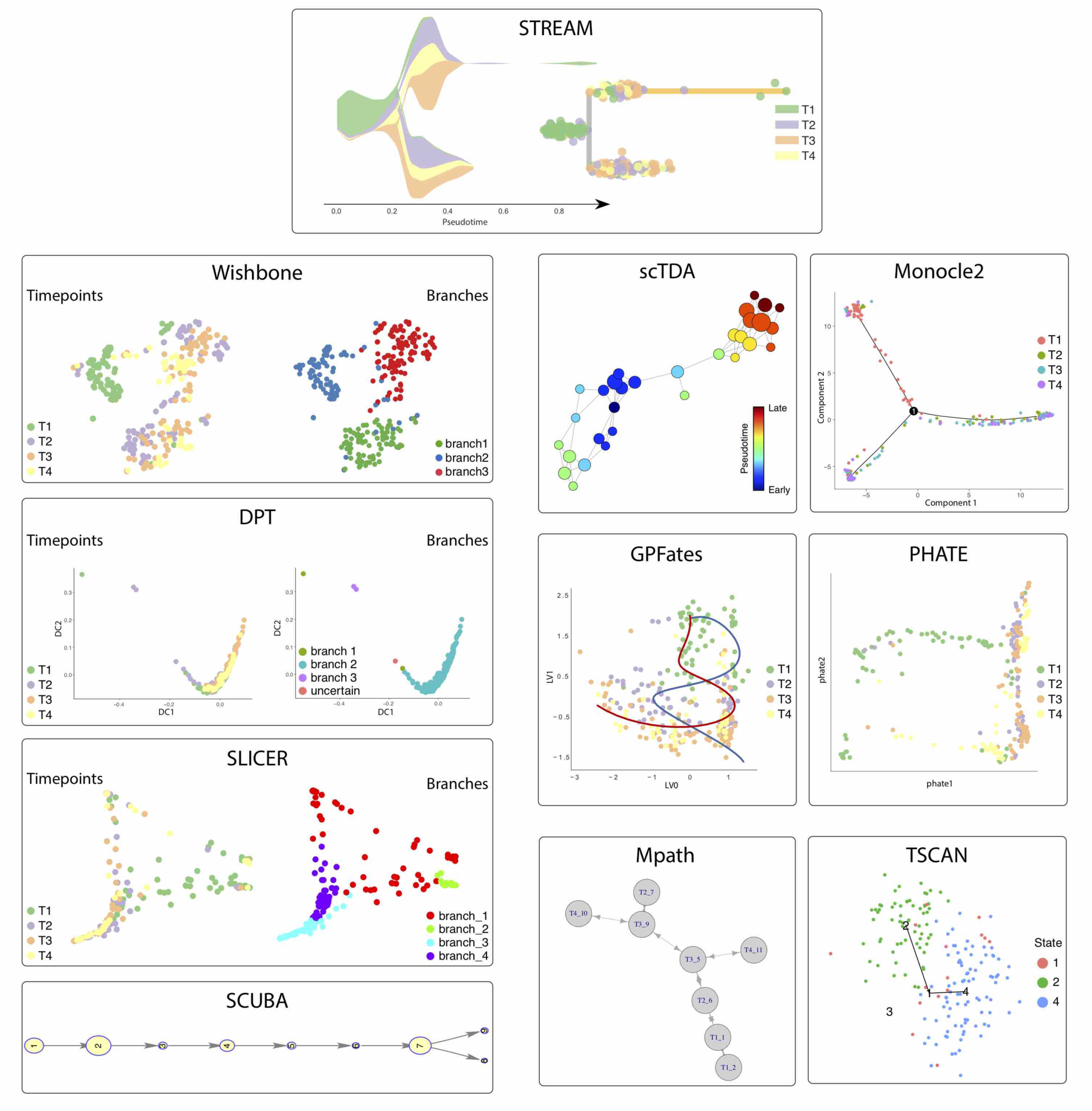
Output of different trajectories inference methods on HSMM scRNA-seq data. Output provided by different methods on HSMM scRNA-seq data, as described in Supplementary Figure 8.

**Supplementary Figure 11.**
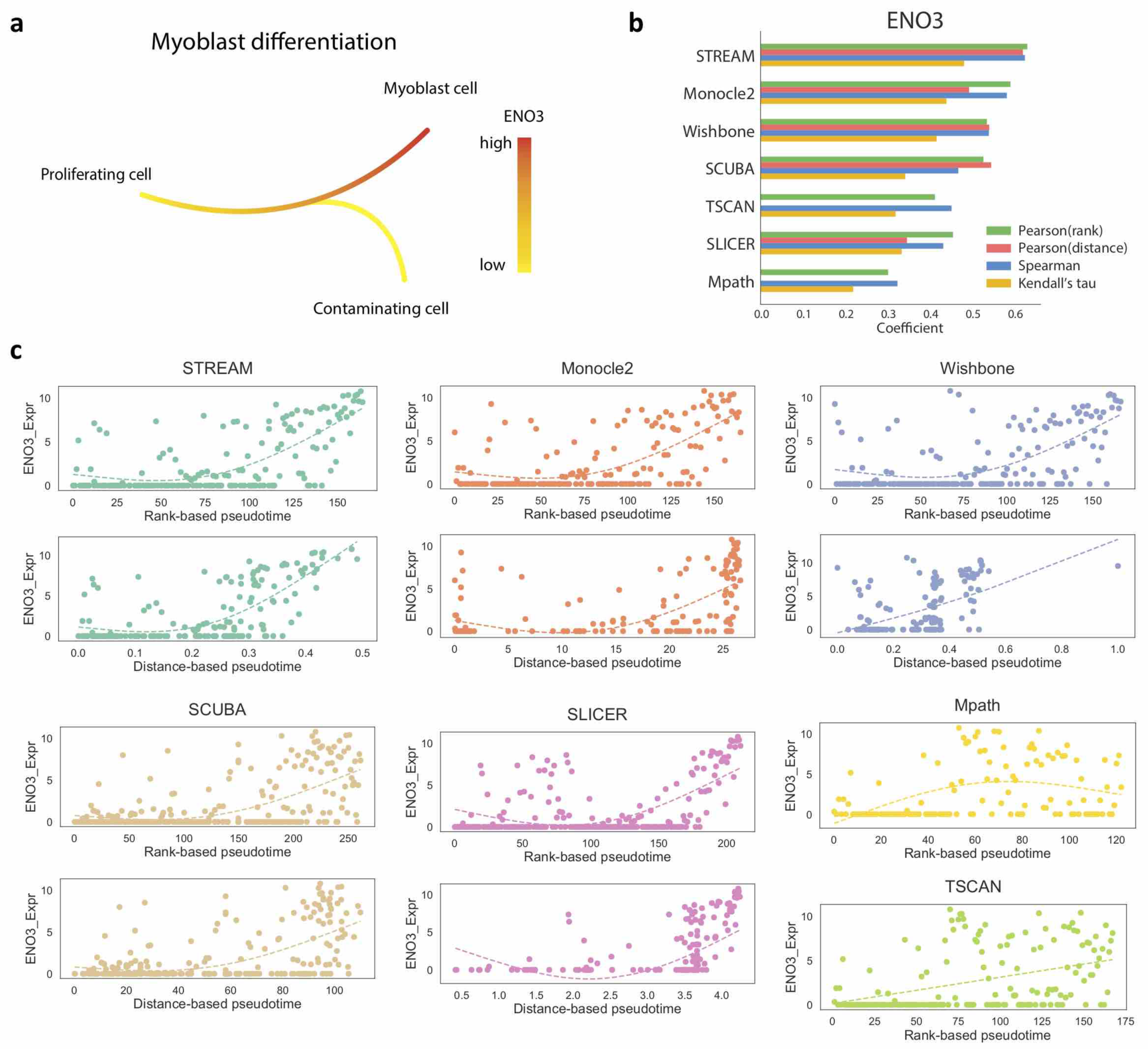
Analysis results of different methods on HSMM scRNA-seq data. (a) ENO3, a marker gene for late-stage differentiation, is used to evaluate the myoblast commitment trajectory (highly expressed in fully differentiated cells) (b) Correlation analysis along myoblast differentiation. The bar plot shows for each method the Pearson correlation between inferred rank-based pseudotime and ENO3 expression, Pearson correlation between inferred distance-based pseudotime and ENO3 expression, Spearman correlation between inferred pseudotime and ENO3 expression and Kendall’s tau correlation between inferred pseudotime and ENO3 expression (c) Along the myoblast differentiation trajectory, the scatter plots show the relationship between ENO3 expression and rank-based pseudotime or distance-based pseudotime inferred by different methods. The dashed lines are fitted curve by generalized additive model (GAM). (Only methods that successfully detect the correct bifurcation are included).

**Supplementary Figure 12.**
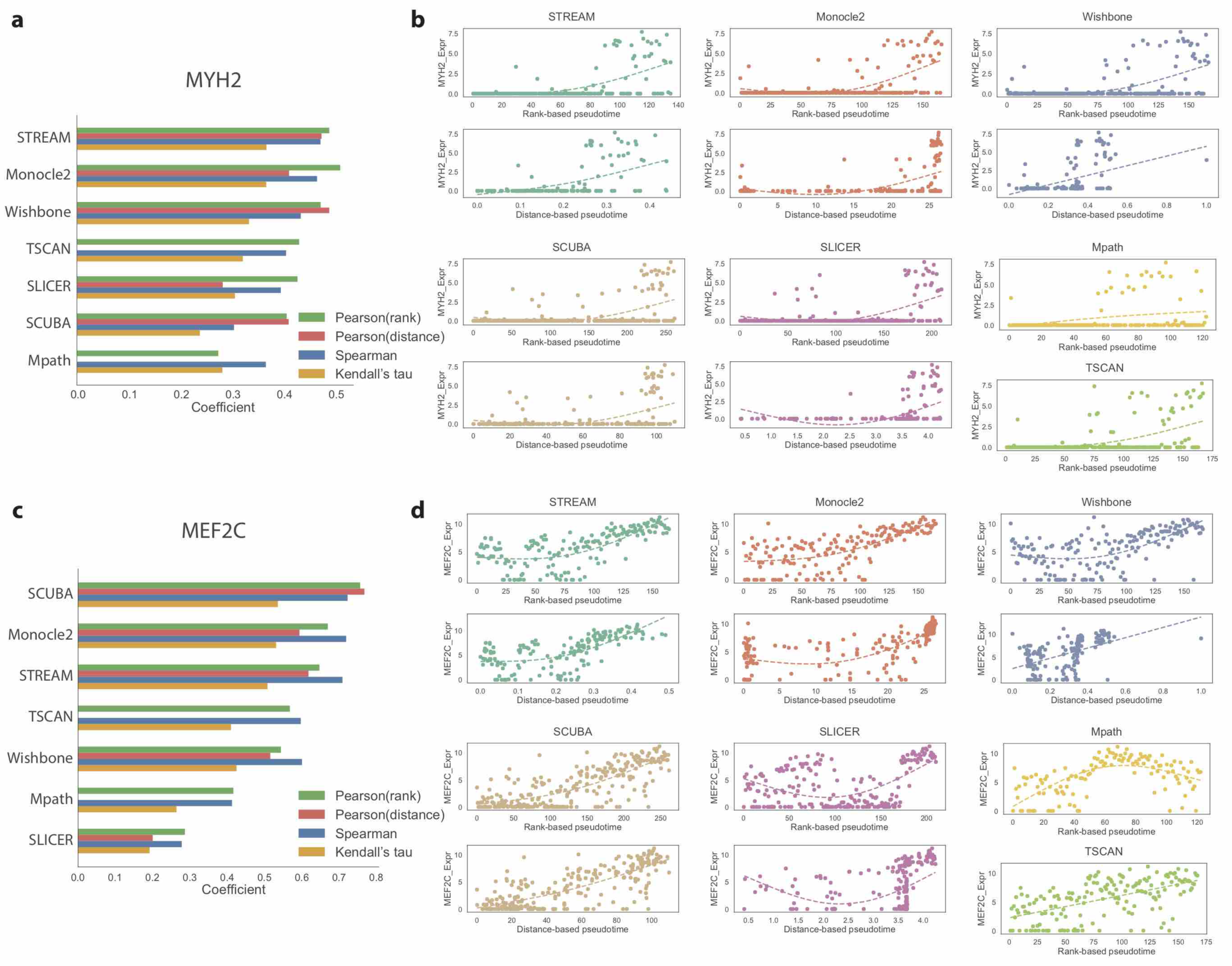
Analysis results of different methods on HSMM scRNA-seq data. Same analysis presented in Supplementary Figure 10 for two additional marker genes of myoblast differentiation: MYH2 and MEF2C

Finally, we analyzed a high-quality single cell qPCR dataset containing ∽270 blood cells sorted from 6 different populations: FISC, MPP, CMP, GMP, MEP and common lymphoid progenitor cells (CLPs) profiled for ∽170 key transcription factors important in mouse hematopoiesis^26^. The output of each method is shown in **Supplementary Fig. 13**. STREAM is the only method that clearly shows the reconstructed developmental trajectories and the lineage hierarchies using its default visualizations. We recovered a trajectory that starts from HSCs and then through MPPs bifurcates into CMPs. A subset of likely erythroid-poised CMPs shows an early progression into MEP, consistent with a recently refined model of hematopoiesis^27^. We also recovered a second bifurcation event that effectively captures cell commitment from MPPs into GMPs and CLPs.

**Supplementary Figure 13.**
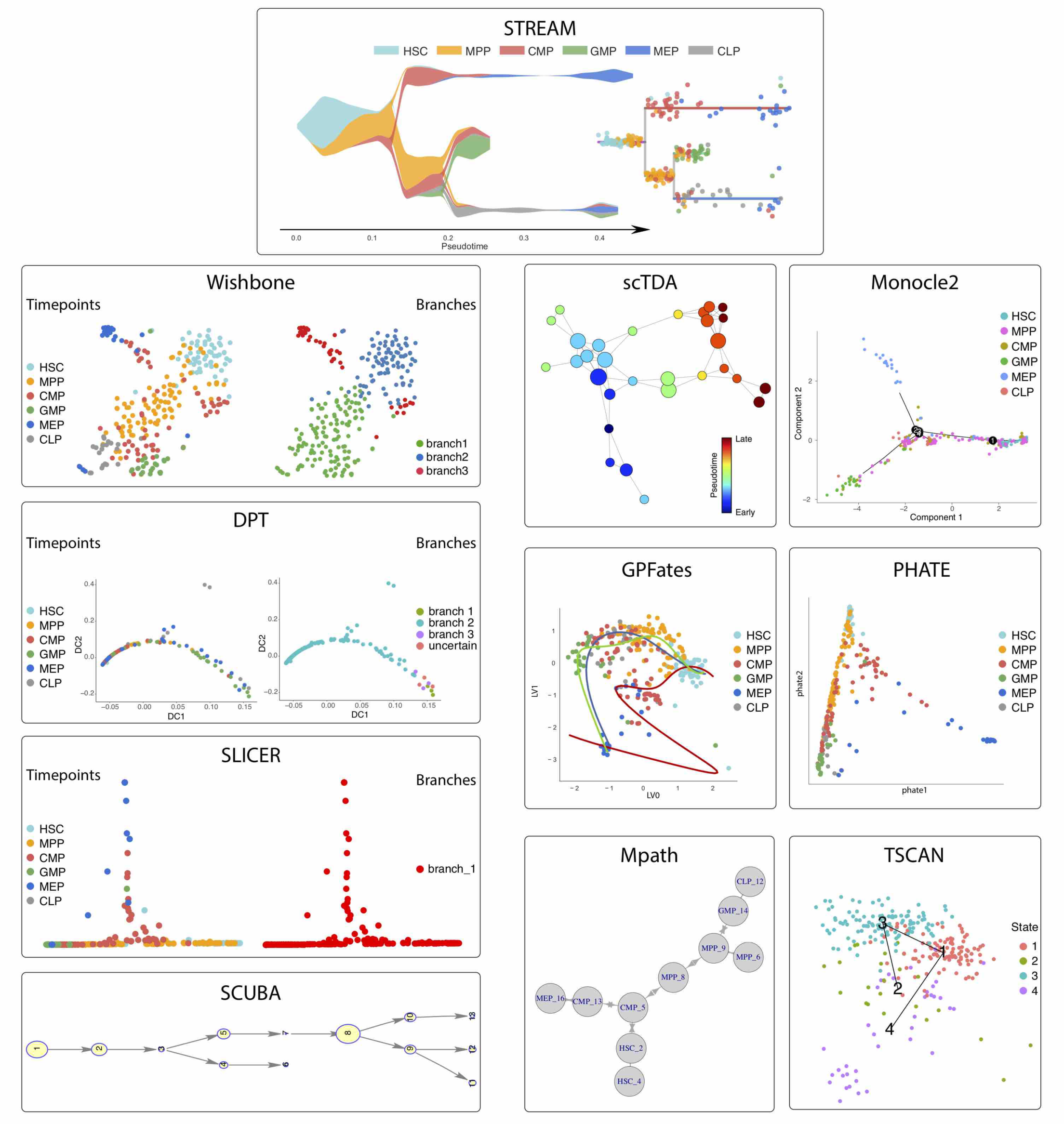
Output of different trajectories inference methods on mouse hematopoietic sc-qPCR data. Output provided by different methods on mouse hematopoietic sc-qPCR data, as described in Supplementary Figure 8.

To assess the quality of the discovered trajectories, we reasoned that classic marker genes for different lineages should be expressed in cells belonging to different trajectories with minimal mixing (i.e. it should be rare to observe single cells that express simultaneously both markers). To this end, we selected *Gata1*, a classic erythroid marker, and *Pax5*, a classic lymphoid marker. For each method, we selected the two best single trajectories that contained *Gata1* or *Pax5* expressing cells (we observed that for some models these two trajectories coincide). Then, each trajectory is evaluated based on precision, recall and the F1 score (see **Online Methods**). The optimal model should balance precision and recall separating *Gata1* and *Pax5* in two distinct trajectories; whereas under-branching models will have a high recall but poor precision and over-branching models will have a high precision but poor recall (**Supplementary Fig. 14a**).

STREAM has the highest F1-score for both *Gata1* and *Pax5* among all the methods tested and balance well precision and recall (**Supplementary Fig. 14b-c**). SCUBA works reasonably well for both genes but has a lower recall overall. Monocle2 tends to generate over-branched structures with high precision but poor recall. Mpath works well in the case of *Gata1* but performs poorly for *Pax5.* Since both Wishbone and DPT can only detect a single bifurcation, they both fail to separate the lymphoid cells from erythroid cells, which leads to having a perfect recall but very poor precision.

In summary, although many of the proposed methods work reasonably well with simple linear trajectories, they may provide over- or under-branched models in more complex scenarios and may mask important trajectories or marker genes.

**Supplementary Figure 14.**
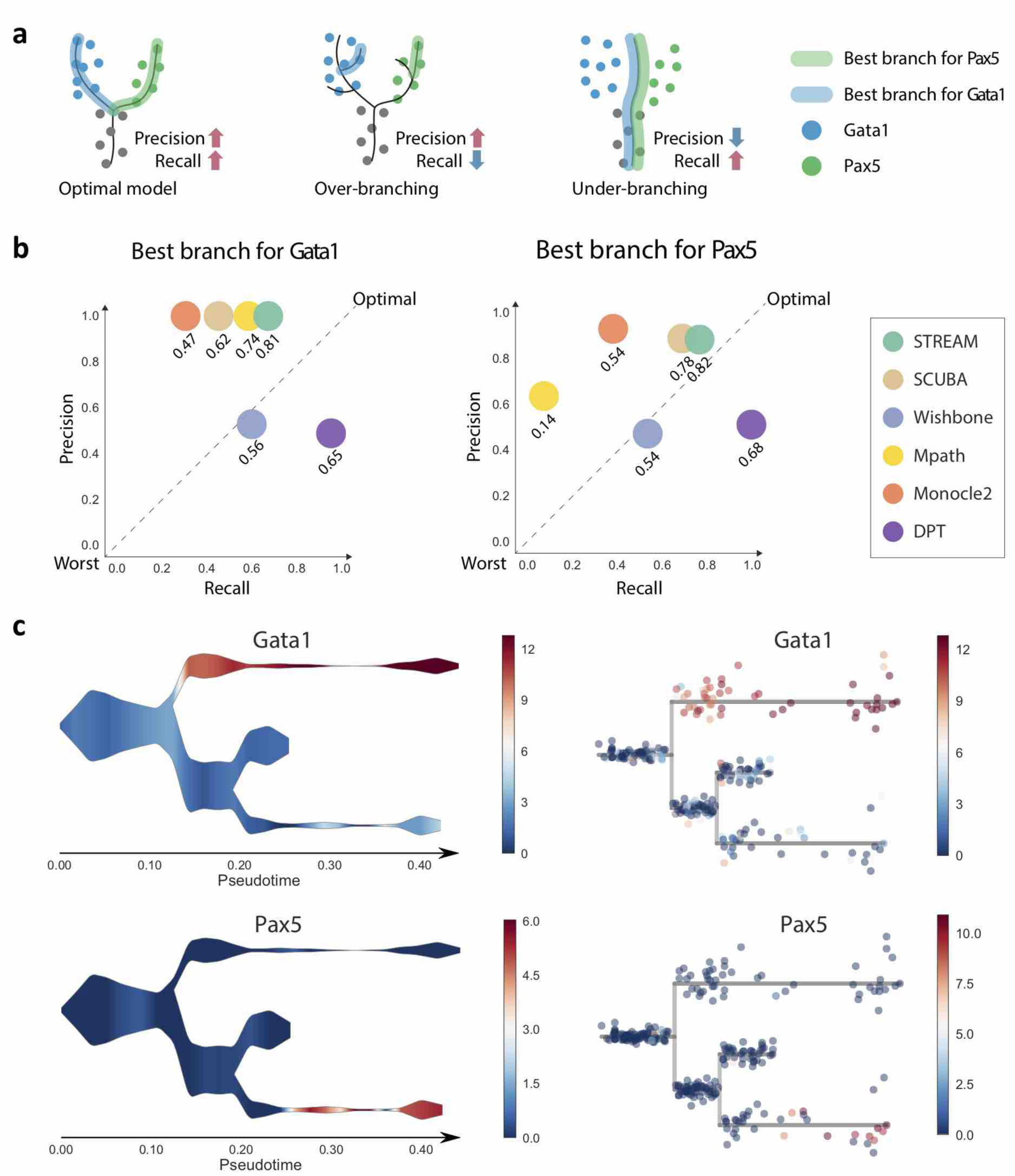
Analysis results of different methods on mouse hematopoietic sc-qPCR data. (a) Using as an example the two marker genes Gata1 and Pax5, which are highly expressed on MEP-committed branch and CLP-committed branch respectively, and rarely co-expresses in the same single cell, the illustration shows three different reconstruction scenarios: optimal model (high precision and high recall), over-branching (high precision but low recall) and under-branching (low precision but high recall). (b) The scatter plots show the relationship between precision and recall for each marker gene across methods. Each circle represents one method and its F_x_ score is reported below. (c) Gata1 and Pax5 expression are visualized in both stream plots and subway map plots for the two best branches automatically detected by STREAM.

### Supplementary Note 6: STREAM analysis on scATAC-seq data of the human hematopoietic system

Although some initial attempts have recently shown how to reconstruct trajectories from single-cell chromatin accessibility data^28,29^, STREAM is the only method to our knowledge that is capable of capturing trajectories from human single-cell (sc) ATAC-seq data using a set of unbiased DNA sequence features. In addition, STREAM is the only end-to-end pipeline that provides specific functions to analyze scATAC-seq data. In fact, Monocle2, a method specifically designed for transcriptomic data and used in the aforementioned initial studies, doesn’t provide in the current implementation (last checked on Apr 13, 2018) any tool for trajectory reconstruction based on scATAC-seq data (http://cole-trapnell-lab.github.io/monocle-release/).

To illustrate STREAM analysis on scATAC-seq data, a total of 3,072 cells were profiled from the human bone marrow and isolated by FACS into nine different cellular populations, including HSC, MPP, CMP, CLP, LMPP, GMP, MEP, mono and plasmacytoid dendritic cells (pDCs)^30^. After filtering cells as previously described,^31^ single cell accessibility profiles for 2,034 high-quality cells passed quality control. We emphasize that each cell was sorted using multiple surface markers as previously described ^32^, providing a phenotypic “true positive” for cell state that would enable us to determine the accuracy of STREAM.

To consider general features related to chromatin accessibility, as many transcription factor motifs have been defined from the hematopoietic system, we sought to determine the efficacy of our approach using general DNA sequence features, i.e. unbiased k-mer scores, which are naive to any known transcription factor motifs and thus generalizable to other systems. To this end, starting from count data and using chromVAR^33^, we first constructed a matrix of cells *x* k-mer accessibility z-scores in our dataset (in our experiments k=7). The k-mer accessibility z-scores are then used by STREAM as features to reconstruct trajectories (**Online Methods**).

STREAM accurately reconstructed cellular developmental trajectories of the human blood system: the HSCs branch segregates through MPP into the erythrocyte-committed, lymphocyte-committed and myelocyte-committed branches. STREAM also reconstructed the bifurcation from lymphoid multipotent progenitors (LMPP) to CLP and plasmacytoid dendritic cells (pDC). Interestingly, STREAM reveals a similar and consistent hematopoietic hierarchy described by orthogonal assays such as transcriptomic profiling (**Fig. 2b**).

**Supplementary Figure 15.**
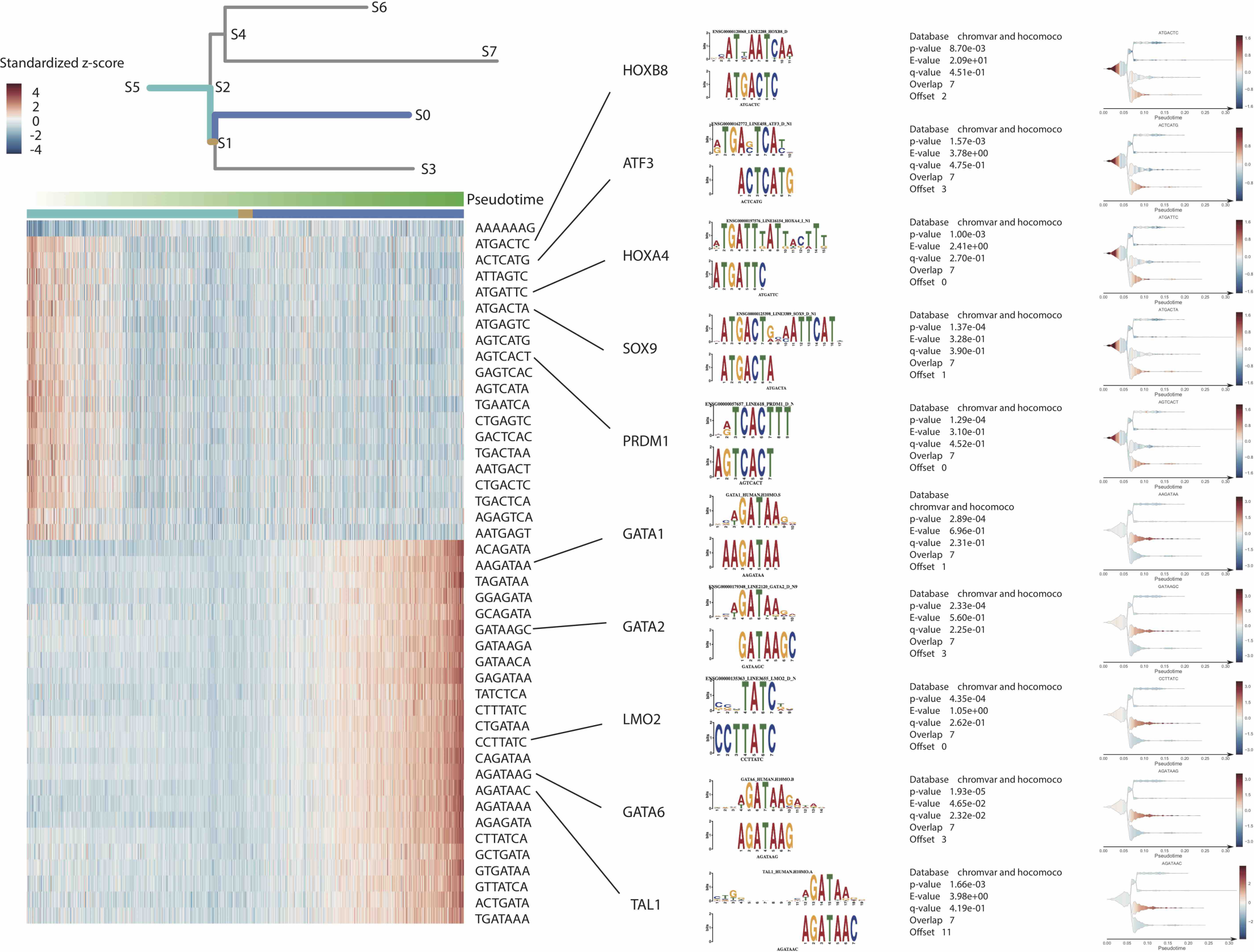
STREAM automatically discovers important k-mers along the erythroid cells differentiation trajectory (S5,S2,S1,S0) for the human hematopoietic system. Top left, subway map schematic to highlight the erythroid cells differentiation trajectory (S5,S2,S1,S0) consisting of three branches (S5,S2)(cyan), (S2,S1)(brown) and (S1,S0)(blue), whose related marker k-mers are calculated. Bottom left, heatmap showing the standardized Z-scores of detected important k-mers. Each row indicates one k-mer. Each column indicates one cell on the trajectory (S5,S2,S1,S0) and cells are ordered by the inferred pseudotime from STREAM. The three colors represent different cell branch ID assignments including branch (S5,S2), branch(S2,S1), and branch(S1,S0). Right, ten detected k-mers selected from left heatmap and their target transcription factor motifs along with output information reported by the motif comparison tool Tomtom. These k-mers are further visualized on stream plots.

In addition, STREAM, using the inferred structure, automatically identifies significant k-mer DNA sequences for each branch. Importantly, those recovered k-mers can be mapped to known transcription factors motifs that may drive cell-fate decision and commitment. In addition to GATA1 and CEPBA, recovered for erythroid lineage and myeloid lineages, respectively (**Fig. 2c**), along the erythroid cell differentiation trajectory, we uncovered several additional potential regulators for HSCs (ATF3^34^, HOXB8^35^) and MEPs (LMO2^36^,TAL1^30^) (**Supplementary Fig.15**).

In summary, compared to previous studies, STREAM provides the unbiased reconstruction of human hematopoiesis using chromatin accessibility data at single-cell resolution. Moreover, it uncovers annotated (i.e. mappable to transcription factors) or unannotated DNA sequences that may be important in defining the different developmental paths.

### Supplementary Note 7: STREAM interactive website

In order to make STREAM user-friendly and accessible to non-bioinformatician, we have created an interactive website: http://stream.pinellolab.org. The website implements all the features of the command line version and provides interactive and exploratory panels to zoom and visualize single-cells on any given branch.

The website offers two functions: 1) to run STREAM on single-cell transcriptomic or epigenomic data provided by the users and 2) the first interactive database of precomputed trajectories with results for seven published datasets. The users can visualize and explore cells’ developmental trajectories, subpopulations and their gene expression patterns at single-cell level. (**Supplementary Fig. 16**).

The website can also run on a local machine using the provided Docker image we have created. To run the website in a local machine it is just necessary to install Docker and then from the command line execute the following command:

docker run -p 10001:10001 pinellolab/stream STREAM_webapp

After the execution of the command the user will have a local instance of the website accessible at the URL: http://localhost:10001

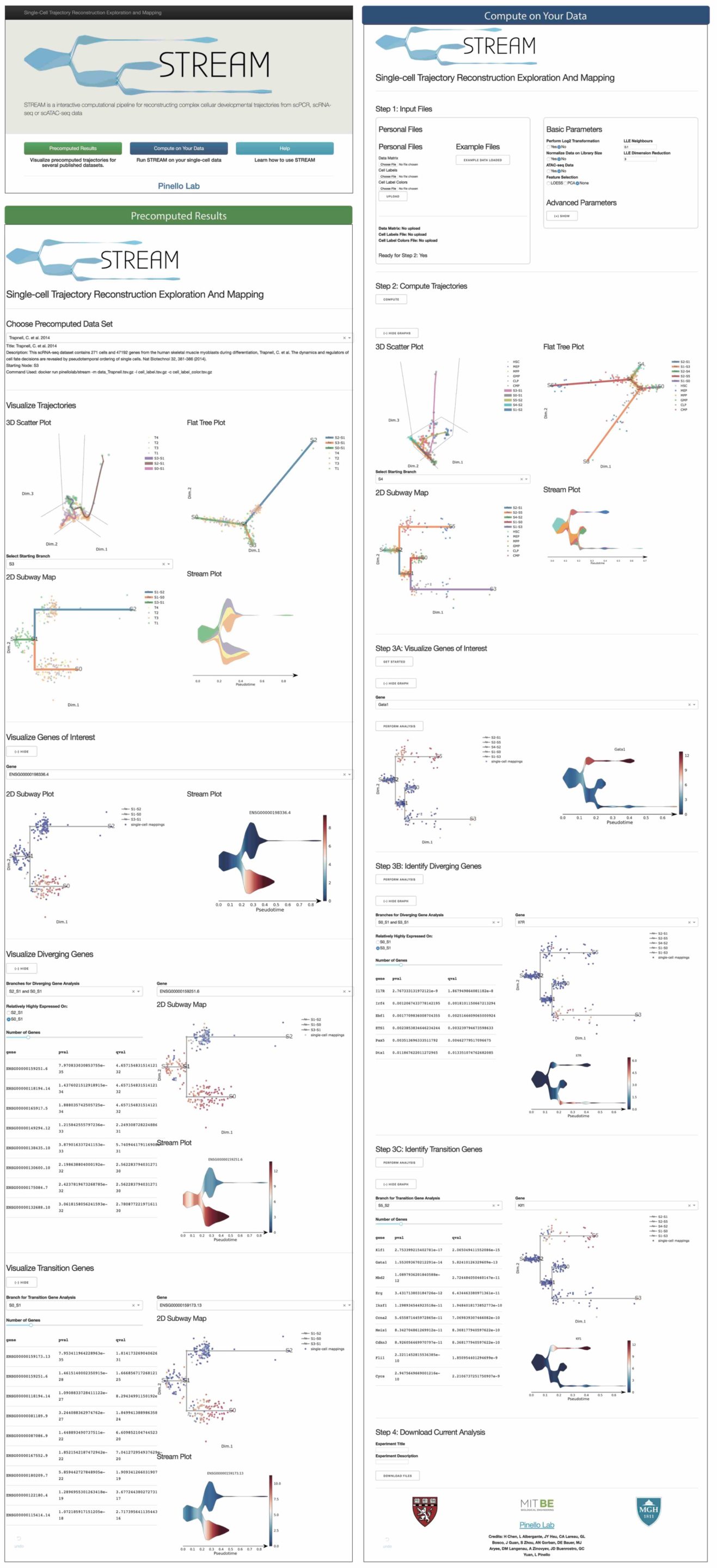
STREAM interactive website. It consists of ‘precomputed results’ and ‘Compute on Your Data’. In ‘Precomputed Results’, users can visualize single cell trajectories from several published datasets. In ‘Compute on Your Data’, in addition to visualization, users can also infer trajectory using their personal dataset.

### Supplementary Note 8: STREAM command line interface

STREAM can be easily used thanks to a simple command line interface. It is possible to install and use STREAM with Docker.

### Installation with Docker

With Docker no installation is required, the only dependence is Docker itself. Docker can be downloaded freely from here: https://store.docker.com/search?offering=community&type=edition

To get a local copy of STREAM execute the following command:
docker pull pinellolab/stream

### STREAM usage and example dataset

The main and required input file is a tab-separated gene expression matrix (raw counts or scaled) in tsv file format. Each row represents a unique gene and each column is one cell.

The following table shows the first 5 rows (genes) and 5 columns (cells) of the provided example dataset

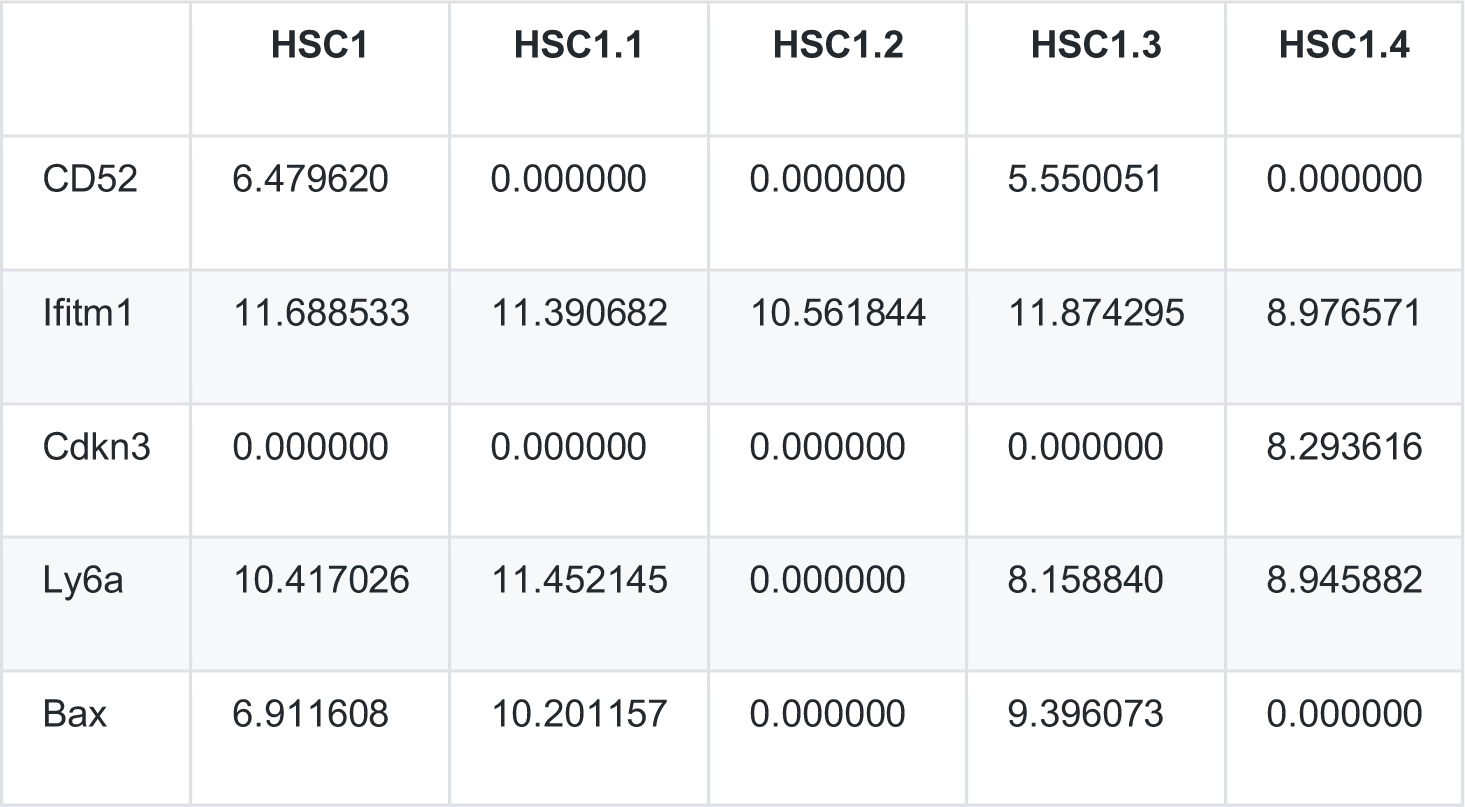

|  | HSC1 | HSC1.1 | HSC1.2 | HSC1.3 | HSC1.4 |
| --- | --- | --- | --- | --- | --- |
| CD52 | 6.479620 | 0.000000 | 0.000000 | 5.550051 | 0.000000 |
| Ifitm1 | 11.688533 | 11.390682 | 10.561844 | 11.874295 | 8.976571 |
| Cdkn3 | 0.000000 | 0.000000 | 0.000000 | 0.000000 | 8.293616 |
| Ly6a | 10.417026 | 11.452145 | 0.000000 | 8.158840 | 8.945882 |
| Bax | 6.911608 | 10.201157 | 0.000000 | 9.396073 | 0.000000 |

In addition, it is possible to provide these optional files in .tsv format:

1. **Cell labels:** Each item can be a putative cell type or sampling time point obtained from experiments. Cell labels are helpful for visually validating the inferred trajectory. This file must be in .tsv format. The order of labels should be consistent with cell order in the gene expression matrix file. No header is necessary:

HSC
HSC
GMP
MEP
MEP
MPP
GMP
GMP
.
.
.
2. **Cell label color:** Customized colors to use for the different cell labels. The first column specifies cell labels and the second column specifies the color in the format of hex. No header is necessary:

HSC #7DD2D9
MPP #FFA500
CMP #e55b54
GMP #5dab5a
MEP #166FD5
CLP #989797
3. **Gene list:** It contains genes that users may be interested in visualizing in subway map and stream plot in addition to the genes detected by STREAM. Genes are listed in one column. No header is necessary:

Ifitm1
Cdkn3
Ly6a
CD52
Foxo1
4. **Feature genes:** It contains genes that the user can specify and that are used as features to infer trajectories. instead of using the automatic feature selection of STREAM. No header is necessary:

Gata1
Pax5
CD63
Klf1
Lmo2
.
.
.

To run STREAM, after the installation at the command-line interface execute:
docker run pinellolab/stream [options]

Users can specify the following options:

~~~
-m, ‐‐matrix
input file name. Matrix is in .tsv or tsv.gz format in which each row represents a unique gene and each column is one cell. (default: None)
-l, ‐‐ cell_labels
file name of cell labels (default: None)
-c, ‐‐cell_labels_colors
file name of cell label colors (default: None)
-s, ‐‐select_features
LOESS, PCA, all: Select variable genes using LOESS or top principal components using PCA or keep all the gene (default: LOESS)
-f, ‐‐feature_genes
specified feature genes (default: None)
-t, ‐‐detect_TG_genes
detect transition genes automatically -d, ‐‐detect_DE_genes detect DE genes automatically -g, ‐‐gene_list
genes to visualize, it can either be filename which contains all the genes in one column or a set of gene names separated by comma (default: None)
-p, ‐‐use_precomputed
use precomputed data files without re-computing structure learning part ‐‐log2
perform log2 transformation ‐‐norm
normalize data based on library size ‐‐atac
indicate scATAC-seq data ‐‐atac_counts
scATAC-seq counts file name in .tsv or .tsv.gz format. Counts file is a compressed sparse matrix that contains three columns including region indices, sample indices and the number of reads(default: None)
‐‐atac_regions
scATAC-seq regions file name in .tsv or .tsv.gz format. Regions file contains three columns including chromosome names, start and end positions of regions (default: None)
‐‐atac_samples
scATAC-seq samples file name in .tsv or tsv.gz. Samples file contains one column of cell names (default: None)
‐‐atac_k
specify k-mers length for scATAC-seq analysis (default: 7)
‐‐n_processes
Specify the number of processes to use. (default, all the available cores).
‐‐loess_frac
The fraction of the data used in LOESS regression (default: 0.1)
‐‐pca_max_PC
Maximal principal components in PCA (default: 100)
‐‐pca_first_PC keep first PC ‐‐pca_n_PC
The number of selected PCs (default: 15)
‐‐n_processes
Specify the number of processes to use. The default uses all the cores available ‐‐lle_neighbours
LLE neighbour percent (default: 0.1)
‐‐lle_components
number of components for LLE space (default: 3)
‐‐AP_damping_factor
Affinity Propagation: damping factor (default: 0.75)
‐‐EPG_n_nodes
Number of nodes for elastic principal graph (default: 50)
‐‐EPG_lambda
lambda parameter used to compute the elastic energy (default: 0.02)
‐‐EPG_mu
mu parameter used to compute the elastic energy (default: 0.1)
‐‐EPG_trimmingradius
maximal distance of point from a node to affect its embedment (default: Inf)
‐‐EPG_finalenergy
indicating the final elastic energy associated with the configuration. It can be ‘Base’ or ‘Penalized’ (default: ‘Penalized’)
‐‐EPG_alpha
positive numeric, alpha parameter of the penalized elastic energy (default: 0.02) ‐‐disable_EPG_collapse
disable collapsing small branches
‐‐EPG_collapse_mode
the mode used to collapse branches. It can be ‘PointNumber’,‘PointNumber_Extrema’, ‘PointNumber_Leaves’,‘EdgesNumber’ or ‘EdgesLength’ (default:‘PointNumber’)
‐‐EPG_collapse_par
the control parameter used for collapsing small branches
‐‐EPG_shift
shift branching point
‐‐EPG_shift_mode
the mode to use to shift the branching points ‘NodePoints’ or ‘NodeDensity’ (default: NodeDensity) ‐‐EPG_shift_DR
positive numeric, the radius used when computing point density if EPG_shift_mode is ‘NodeDensity’ (default:0.05)
‐‐EPG_shift_maxshift
positive integer, the maximum distance (number of edges) to consider when exploring the branching point neighborhood (default:5)
‐‐disable_EPG_ext
disable extending leaves with additional nodes ‐‐EPG_ext_mode
the mode used to extend the graph. It can be ‘QuantDists’, ‘QuantCentroid’ or ‘WeigthedCentroid'. (default: QuantDists)
‐‐EPG_ext_par
the control parameter used for contribution of the different data points when extending leaves with nodes (default: 0.5)
‐‐DE_z_score_cutoff
Differentially Expressed Genes Z-score cutoff (default: 2)
‐‐DE_diff_cutoff
Differentially Expressed Genes difference cutoff (default: 0.2)
‐‐TG_spearman_cutoff
Transition Genes Spearman correlation cutoff (default: 0.4)
‐‐TG_diff_cutoff
Transition Genes difference cutoff (default: 0.2)
‐‐stream_log_view
use log2 scale for y axis of stream_plot ‐‐for_web
Output files for website
-o, ‐‐output_folder
Output folder (default: None)
‐‐new
file name of data to be mapped (default: None)
‐‐new_l
filename of new cell labels (default: None)
‐‐new_c
filename of new cell label colors (default: None)
~~~

***Example with transcriptomic data:*** Using the example data provided: data_guoji.tsv, cell_label.tsv and cell_label_color.tsv, and assuming that they are in the current folder, to perform trajectories analysis, users can simply run a single command (By default, LOESS is used to select most variable gene. For qPCR data, the number of genes is relatively small and often preselected, it this case it may be necessary to keep all the genes as features by setting the flag-s all):
docker run pinellolab/stream -v $PWD:/data -w /data -m data_guoji.tsv -l cell_label.tsv -c cell_label_color.tsv -s all

To visualize genes of interest, user can provide a gene list file, for example: gene_list.tsv and add the flag -p to use the precomputed file obtained from the first running (in this way, the analysis can will not restart from the beginning and other existing figures will not be re-generated):
docker run pinellolab/stream -v $PWD:/data -w /data -m data_guoji.tsv -l cell_label.tsv -c cell_label_color.tsv -s all -g gene_list.tsv -p

To explore potential marker genes, it is possible to add the flags -d or -t to detect DE (differentially expressed) genes and transition gens respectively. The best 10 DE (any pair of branches) and transition genes (any branch) are automatically plotted:
docker run pinellolab/stream -v $PWD:/data -w /data -m data_guoji.tsv -l cell_label.tsv -c cell_label_color.tsv -s all -d -t

***Example of the mapping feature:*** To use the mapping feature, users need to provide two datasets: one used to inferring trajectories and the other containing cells to be mapped to the inferred trajectories. Here to illustrate this feature we use data from Moore et al.2016 and in particular those two files: Moore.tsv (cells used to infer trajectories), data_mapping.tsv (cells to map).

We first infer trajetories using the following command:
docker run pinellolab/stream -v $PWD:/data -w /data -m data_Moore.tsv -s all ‐‐ EPG_shift ‐‐EPG_trimmingradius 0.1 -o /users_path/STREAM_result

Then to map the cells to the inferred trajectories we use the following command (note the flags ‐‐new, ‐‐new_l and ‐‐ new_c and the same folder for the output flag -o):
docker run pinellolab/stream -v $PWD:/data -w /data -o /users_path/STREAM_result ‐‐new data_mapping.tsv ‐‐new_l cell_labels_mapping.tsv ‐‐new_c cell_labels_mapping_color.tsv

After running this command, a folder named Mapping_Result will be created under /users_path/STREAM_result along with all the mapping analysis results.

***Example with scATAC-seq data:*** To perform scATAC-seq trajectory inference analysis, three files are necessary, a .tsv file of counts in compressed sparse format, a sample file in .tsv format and a region file in .bed format:

1. **Count file:** a tab-delimited compressed matrix in sparse format (column-oriented). It contains three columns. The first column specifies the rows indices (the regions) for non-zero entry. The second column specifies the columns indices (the sample) for non-zero entry. The last column contains the number of reads in a given region for a particular cell. No header is necessary:

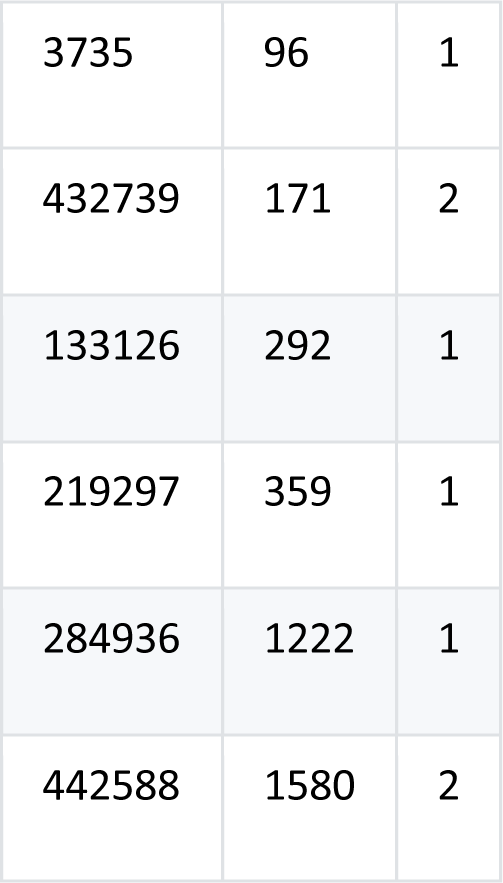
2. Sample file: It has one column. Each row is a cell name. The order of the cells should be consistent with the sample indices in count file. No header is necessary:

singles-BM0828-HSC-fresh-151027-1
singles-BM0828-HSC-fresh-151027-2
singles-BM0828-HSC-fresh-151027-3
singles-BM0828-HSC-fresh-151027-4
singles-BM0828-HSC-fresh-151027-5
3. **Region file:** a tab-delimited .bed file with three columns. The first column specifies chromosome names. The second column specifies the start position of the region. The third column specifies the end position of the region. The order of regions should be consistent with the regions indices in the count file. No header is necessary:

chr1 10279 10779
chr1 13252 13752
chr1 16019 16519
chr1 29026 29526
chr1 96364 96864
.
.
.

Using these three files, users can run STREAM with the following command (note the flag ‐‐atac):
docker run pinellolab/stream -v $PWD:/data -w /data ‐‐atac -s PCA ‐‐atac_counts count_file.tsv ‐‐atac_sample_file.tsv ‐‐atac_regions region_file.bed -l cell_label.tsv -c cell_label_color.tsv

This command will generate a file named df_zscores_scaled. tsv. It’s a tab-delimited z-score matrix with k-mers in row and cells in column. Each entry is a scaled z-score of the accessibility of each k-mer across cells. This operation is time-consuming and it may take a couple of hours with a modest machine. STREAM also provides the option to take as input a precomputed z-score file from the previous step, for example, to recover trajectories when increasing the dimensionality of the manifold. Using a precomputed z-score file, users can run STREAM with the following command:
docker run pinellolab/stream -v $PWD:/data -w /data -m df_zscores_scaled.tsv -l cell_label.tsv -c cell_label_color.tsv ‐‐atac -s PCA

#### Output description

STREAM write all the results by default in the folder STREAM_result, unless a different directory is specified by the user with the flag -o. This folder contains the following files and directories:

- **LLE.pdf:** projected cells in the MLLE 3D space.
- **EPG.pdf:** elastic principal graph fitted by ElPiGraph in 3D space
- **flat_tree.pdf:** 2D single-cell level flat tree plot
- **nodes.tsv:** positions of nodes (or states) in the flat_tree plot
- **edges.tsv:** edges information in the flat_tree plot
- **cell_info.tsv:** Cell information file. Column ‘CELL_ID’, the cell names in the input file. Column ‘Branch’, the branch id a cell is assigned to. The branch id is encoded by the two cell states. Column ‘lam’, the location on a branch, which is the arc length from the first cell state of branch id to the projection of the cell on that branch. Column ‘dist’, the euclidian distance between the cell and its projection on the branch.
- sub-folder **‘Transition_Genes’** contains several files, one for each branch id, for example for (S1,S2):

- **Transition_Genes_S1_S2.png:** Detected transition genes plot for branch S1_S2. Orange bars are genes whose expression values increase from state S1 to S2 and green bars are genes whose expression values decrease from S1 to S2
- **Transition_Genes_S1_S2.tsv:** Table that stores information of detected transition genes for branch S1_S2.
- sub-folder **‘DE_Genes’ Genes’** contains several files, one for each pair of branches, for example for (S1,S2) and (S3,S2):

- **DE_genes_S1_S2 and S3_S2.png:** Detected differentially expressed top 15 genes plot. Red bars are genes that have higher gene expression in branch S1_S2, blue bars are genes that have higher gene expression in branch S3_S2
- **DE_up_genes_S1_S2 and S3_S2.tsv:** Table that stores information of DE genes that have higher expression in branch S1_S2.
- **DE_down_genes_S1_S2 and S3_S2.tsv:** Table that stores information of DE genes that have higher expression in branch S3_S2.
- sub-folder **‘S0’:** Set of linearized plots (subway and stream plots) for each of the cell states, for example, choosing S0 state as root state:

- **subway_map.png:** single-cell level cellular branches plot
- **stream_plot.png:** density level cellular branches plot
- **subway_map_gene.png:** gene expression pattern on subway map plot
- **stream_plot_gene.png:** gene expression pattern on stream plot
- sub-folder**‘Precomputed‘:**

- It contains files that store computed variables used when the flag -p is enabled.

